# A cellular entity retaining only its replicative core: Hidden archaeal lineage with an ultra-reduced genome

**DOI:** 10.1101/2025.05.02.651781

**Authors:** Ryo Harada, Yuki Nishimura, Mami Nomura, Akinori Yabuki, Kogiku Shiba, Kazuo Inaba, Yuji Inagaki, Takuro Nakayama

## Abstract

Defining the minimal genetic requirements for cellular life remains a fundamental question in biology. Genomic exploration continually reveals novel microbial lineages, often exhibiting extreme genome reduction, particularly within symbiotic relationships. Here, we report the discovery of *Candidatus* Sukunaarchaeum mirabile, a novel archaeon with an unprecedentedly small genome of only 238 kbp —less than half the size of the smallest previously known archaeal genome— from a dinoflagellate-associated microbial community. Phylogenetic analyses place Sukunaarchaeum as a deeply branching lineage within the tree of Archaea, representing a novel major branch distinct from established phyla. Environmental sequence data indicate that sequences closely related to Sukunaarchaeum form a diverse and previously overlooked clade in microbial surveys. Its genome is profoundly stripped-down, lacking virtually all recognizable metabolic pathways, and primarily encoding the machinery for its replicative core: DNA replication, transcription, and translation. This suggests an unprecedented level of metabolic dependence on a host, a condition that challenges the functional distinctions between minimal cellular life and viruses. The discovery of Sukunaarchaeum pushes the conventional boundaries of cellular life and highlights the vast unexplored biological novelty within microbial interactions, suggesting that further exploration of symbiotic systems may reveal even more extraordinary life forms, reshaping our understanding of cellular evolution.

**Graphical Abstract:** 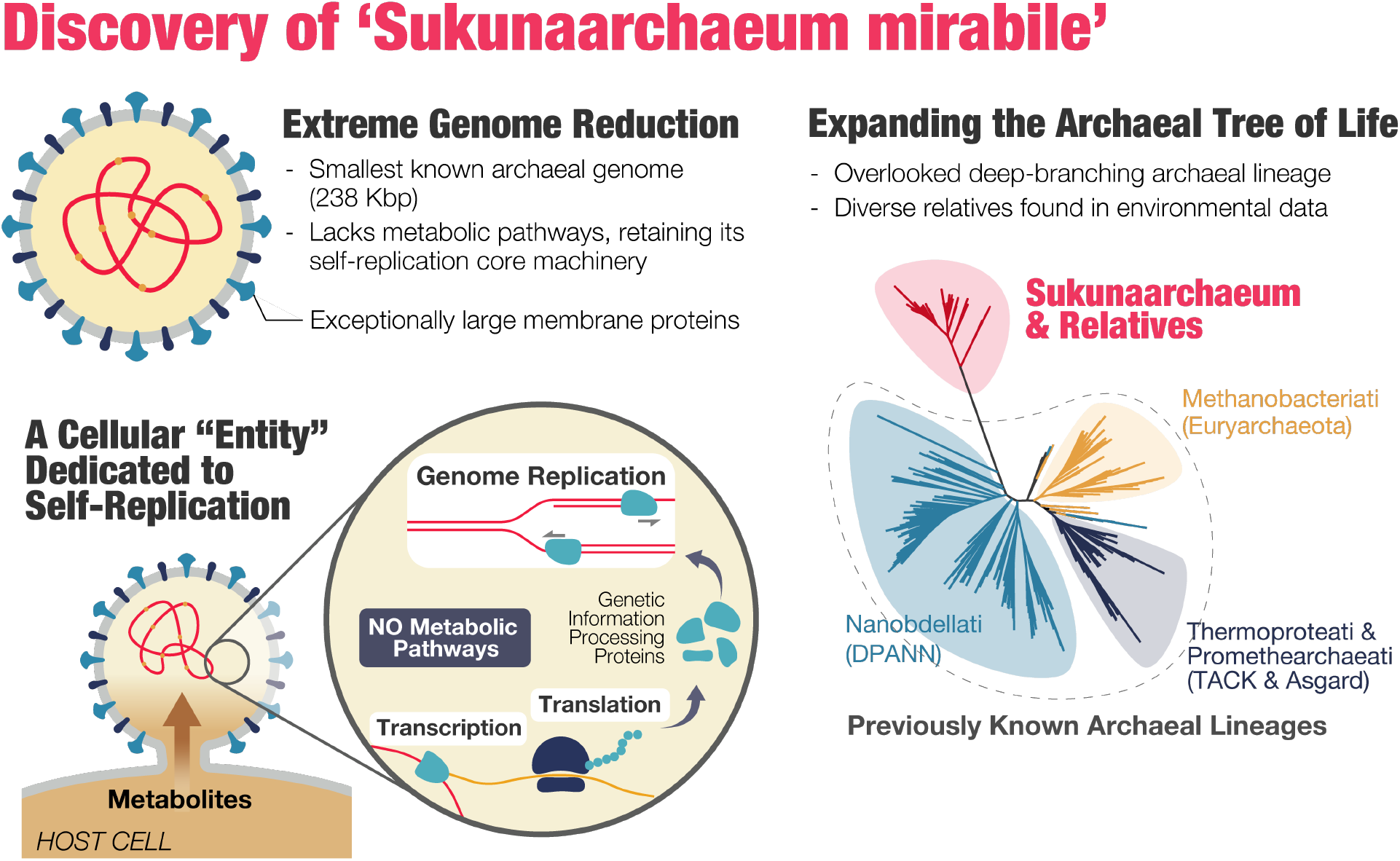

## Introduction

Metagenomic approaches have revolutionized our understanding of microbial diversity, revealing vast numbers of unculturable microorganisms (Hug et al., 2016; Zhu et al., 2019). These discoveries include entirely new phylogenetic groups at the kingdom or phylum level. Examples include Minisyncoccota (also known as the Candidate Phyla Radiation/CPR) within the bacterial domain, as well as Promethearchaeati (also known as Asgard archaea) and Nanobdellati (also known as DPANN) within the archaeal domain. These discoveries have remarkably revised our understanding of biological evolution. The discovery of these novel microbial lineages has also highlighted the crucial role of symbiotic relationships in shaping microbial ecosystems. Minisyncoccota (CPR) bacteria and Nanobdellati (DPANN) archaea, despite belonging to different biological domains, are often compared due to their shared characteristics: having rapid rates of sequence evolution, possessing small genomes, and exhibiting reduced metabolic capacities (Castelle et al., 2018), often leading to obligate symbiotic lifestyles reliant on host resources. Reflecting this presumed dependency, successfully cultured representatives from these groups have consistently shown obligate symbiotic or co-culture-dependent lifestyles (Cross et al., 2019; Golyshina et al., 2017; Hamm et al., 2019, 2024; He et al., 2015; Huber et al., 2002; Kato et al., 2022; Krause et al., 2022; La Cono et al., 2020; Sakai et al., 2022; St John et al., 2019; Wurch et al., 2016). Furthermore, cultured strains of *Prometheoarchaeum syntrophicum*, a member of Promethearchaeati (Asgard archaea), the archaeal group considered closest to eukaryotes (Eme et al., 2023; Rodrigues-Oliveira et al., 2023; Spang et al., 2015; Zaremba-Niedzwiedzka et al., 2017), establish syntrophic relationships with sulfate-reducing bacteria and methanogenic archaea (Imachi et al., 2025, 2020), further demonstrating the metabolic interdependencies within microbial communities. These findings suggest that a significant reservoir of unknown microbial diversity may be concealed within symbiotic interactions.

Symbiotic relationships between prokaryotes are often the focus of attention (Golyshina et al., 2017; Hamm et al., 2019, 2024; He et al., 2015; Huber et al., 2002; Kato et al., 2022; Krause et al., 2022; Kuroda et al., 2022; La Cono et al., 2020; Reva et al., 2023; St John et al., 2019; Wrede et al., 2012; Wurch et al., 2016). Aquatic environments, where many prokaryotes reside, are also abundant in unicellular eukaryotes (Bar-On & Milo, 2019). Recent discoveries highlight the potential for prokaryotic lineages to reside within or on the surface of unicellular eukaryotes (Husnik et al., 2021; Nakayama et al., 2019), as well as new organelles derived from intracellularly symbiotic prokaryotes, such as the nitroplast of *Braarudosphaera bigelowii* and the chromatophore of *Paulinella* (Coale et al., 2024; Sørensen et al., 2024). This suggests that the intracellular and cell-surface environments of unicellular eukaryotes represent a largely unexplored frontier for microbial discovery.

Here, we report the discovery of a novel archaeon, *Candidatus* Sukunaarchaeum mirabile, from a singlecell amplified genome obtained from a dinoflagellate cell harboring a microbial community. This archaeon possesses an extraordinarily small genome (238 kbp) and represents a deeply branching, previously unknown lineage distinct from established archaeal phyla. Its profoundly stripped-down genome, largely devoid of metabolic functions, retains primarily the core machinery for self-replication (DNA replication, transcription, translation). This extreme specialization in genetic propagation challenges our fundamental understanding of the minimal requirements for cellular life by presenting a viable cell seemingly stripped down to its replicative core.

## Results and Discussion

### Discovery and Genomic Characterization of Sukunaarchaeum

Some marine dinoflagellates are known to maintain symbiotic microbial communities. We recently performed single-cell genome amplification of bacteria associated with the dinoflagellate *Citharistes regius* (Nakayama et al., 2024). This genome amplification product revealed genomes of cyanobacterium (the primary symbiont), two gammaproteobacteria (Nakayama et al., 2025), and a highly unusual circular sequence. This 238 kbp sequence has a low GC content (28.9%) (Figure 1). Multiple independent genome assemblies using short-read, long-read, and hybrid approaches recovered this sequence as a gap-free, circular molecule (Supplementary Table S1). Despite its compactness, the genome encodes both large and small subunit rRNA and 31 tRNAs. Based on the quality standards for genomes assembled from environmental sequences (Bowers et al., 2017), the contiguous, single circular nature of this genome, coupled with the coverage of essential RNA genes, provides compelling evidence that this sequence represents the complete genome of a prokaryote. Annotation revealed 189 open reading frames (ORFs), including archaeal marker genes such as elongation factor 1-alpha (EF-1A) (Supplementary Table S2), confirming that the organism harboring this genome is an archaeon. To our knowledge, the smallest known archaeal complete genome is the 490 Kbp genome of ‘*Nanoarchaeum equitans*’ (Kellner et al., 2018; Waters et al., 2003), making the genome discovered in this study less than half that size. Despite its extraordinarily small size, the CheckM2 assessment suggests high completeness of the genome (CheckM2 estimated completeness and contamination scores: 83.68% and 0.18%, respectively). While we cannot definitively exclude the presence of additional genetic elements (e.g., plasmids) besides the discovered circular chromosome, we found no evidence of other archaeal sequences in the amplified genome (see Supplementary Table S3). To further address the possibility that the circular chromosome represents a fragment of a larger genome, we *in silico* fragmented genomes of other small-genomed archaea into ∼240 kbp segments (matching the size of the circular contig) and assessed their completeness using CheckM2. None of these fragments reached 70% completeness, clearly lower than that of the circular genome (Supplementary Figure S1). This further supports the conclusion that the circular genome contig represents the complete archaeon genome. We named this archaeon *Candidatus* Sukunaarchaeum mirabile; the genus name, *Sukunaarchaeum*, is derived from Sukuna-biko-na, a deity of small stature in Japanese mythology. The predicted proteins of *Candidatus* Sukunaarchaeum mirabile (hereafter referred to as Sukunaarchaeum) exhibit significant sequence divergence from other archaea, suggesting a rapid evolutionary rate. While some proteins lacked detectable sequence homologs in similarity searches on primary sequences, structural modeling with AlphaFold2 allowed for functional inference (Supplementary Figure S2). This suggests that Sukunaarchaeum proteins maintain conserved structural folds despite substantial sequence divergence.

**Figure 1.**
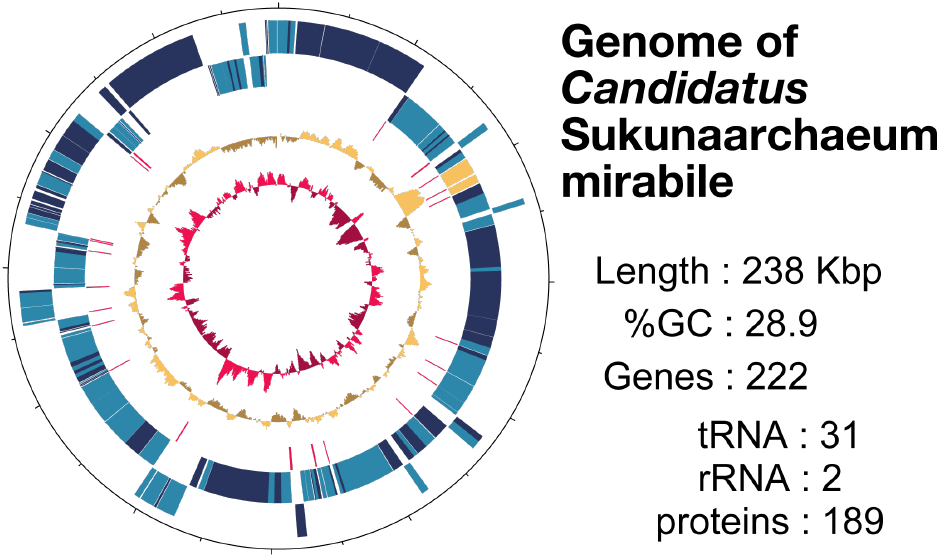
The genome map of Sukunaarchaeum. From outermost to innermost circle, the positions of protein-coding genes and rRNA genes on the +/-strands, tRNA genes, GC content, and GC skew are shown. Color codes for the outermost and 2nd outermost circle: Blue, genes of unknown function; light blue, genes of known function; yellow, rRNA genes.

### Etymology

*Candidatus* Sukunaarchaeum (Su.ku.na.ar.chae’um). N.L. neut. n. Sukuna, derived from Sukuna-biko-na, a deity of small stature in Japanese mythology (treated as a Neo-Latin neuter noun); N.L. neut. n. archaeum (from Gr. adj. archaios, ancient), archaeon; N.L. neut. n. Sukunaarchaeum, an archaeon named after Sukuna-biko-na, referring to the organism’s extremely small genome size. The species name, *mirabile* (mi.ra’bi.le) is derived from L. neut. adj. *mirabile*, meaning marvelous or extraordinary, referring to the remarkable features of this organism, including its unprecedentedly small genome size for an archaeon, its fast sequence evolving rate, and its highly reduced metabolic capabilities.

### Phylogenetic Position of Sukunaarchaeum: A Deep-Branching Archaeal Lineage

We performed comprehensive phylogenetic analyses using a concatenated alignment of 70 conserved archaeal marker proteins from 149 representative archaeal genomes, encompassing all major known archaeal kingdoms and phyla (Parks et al., 2022). We carried out both maximum likelihood (ML) and Bayesian inference (BI) analyses, employing complex substitution models that account for both profile and rate heterogeneity across the sites to accommodate complex evolutionary processes (Baños et al., 2024; Quang et al., 2008). Both ML and BI analyses recovered the monophyly of major archaeal clades, including Nanobdellati (DPANN), Halobacteriota, Thermoplasmatota, Promethearchaeati (Asgardarchaeota) and Thermoproteati (TACK) (Figure 2a; Supplementary Figures S3 and S4), as shown in previous studies (Tahon et al., 2021). Both phylogenetic trees indicated that Sukunaarchaeum branches from a deep position within the archaeal phylogeny and possesses an extremely long branch, over 2.8 times longer than the branch leading to ‘Huberarchaeota’ from the basal node of Nanobdellati clade (6.66 vs. 2.32 substitutions/site), the longest branch within the clade. (Figure 2a). However, the branching position of Sukunaarchaeum differed between the ML and BI methods. In the ML tree, Sukunaarchaeum was positioned as a sister lineage to the entire Nanobdellati clade (Figure 2a; Supplementary Figure S3), whereas in the BI tree, it appeared as a sister lineage to Halobacteriota, one of the phyla comprising Methanobacteriati (formerly known as Euryarchaeota; Supplementary Figure S4). Notably, both branching positions for Sukunaarchaeum had low support values (ultrafast bootstrap value (UFBP) 17 for the ML topology and Bayesian posterior probability (BPP) 0.65 for the BI topology).

**Figure 2.**
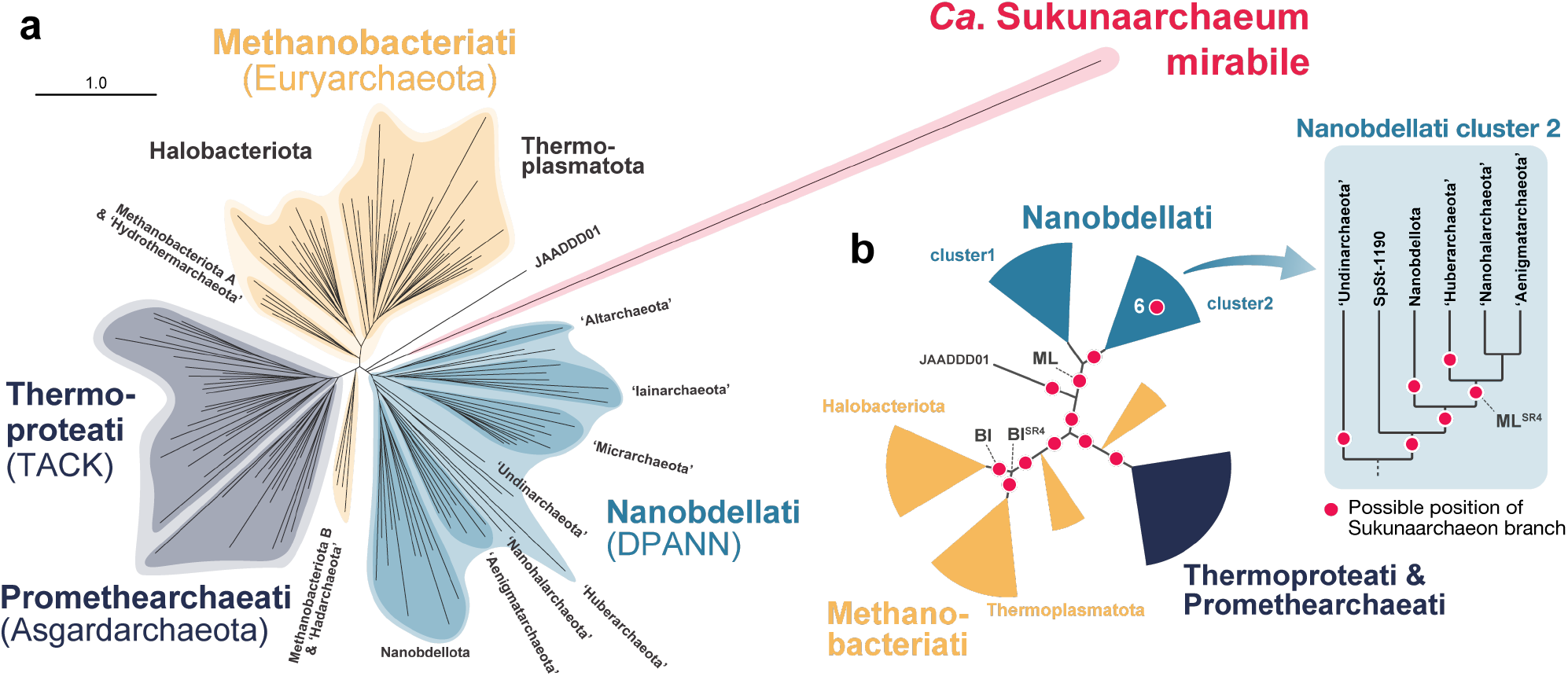
Phylogenetic placement of Sukunaarchaeum within the Archaeal domain. **a**, Maximum likelihood (ML) phylogenetic tree based on a concatenated alignment of 70 conserved archaeal marker proteins. The tree was inferred under the LG+C60+F+I+R10 model, based on a dataset of 150 taxa and 18,286 sites. The scale bar represents the estimated number of substitutions per site. **b**, Exploration of alternative evolutionary positions for Sukunaarchaeum. Red solid circles denote alternative positions for Sukunaarchaeum that were not rejected by the AU test. Nodes labelled as ML and BI represent positions of the Sukunaarchaeum branch in the original trees inferred by ML and BI methods, while ML^SR4^ and BI^SR4^ represent that in the ML and BI tree using the SR4 recoded dataset.

Given the inconsistency between the ML and BI trees and the low support values for the position of Sukunaarchaeum, we performed an approximately unbiased (AU) test to scrutinize all possible branching positions for the Sukunaarchaeum branch within the ML tree topology. With the AU test, while the vast majority of the alternative tree topologies (279 out of 294) were statistically rejected, 15 branching positions, including the one supported by the BI tree, were not. This result suggests 16 possible phylogenetic positions for Sukunaarchaeum (Figure 2b; Supplementary Table S4). These 16 alternative branching positions are all located at the basal positions of the archaeal phylogeny, with some indicating a sister relationship with phyla included in Nanobdellati or Methanobacteriati. Crucially, none of these alternative topologies placed Sukunaarchaeum within any of the known archaeal phyla with statistical confidence.

The branch leading to Sukunaarchaeum is extremely long, more than twice as long as any branch within each of the kingdom-level clades, reflecting the rapid evolutionary rate inferred from its protein sequences. In the context of molecular phylogenetic analysis, there is a known methodological artifact called long-branch attraction (LBA), where lineages with long branches (i.e., fast evolutionary rates) can be artifactually grouped during the tree search process (Felsenstein, 1978; Wang et al., 2008). As a result, they may be falsely positioned in a way that differs from their true phylogenetic positions. It is possible that the Sukunaarchaeum branch is influenced by LBA, given its exceptionally long branch. Furthermore, the Sukunaarchaeum genome exhibits an exceptionally low GC content (28.9%), indicating a skewed nucleotide composition that could influence amino acid frequencies in the encoded proteins. Such compositional biases are also known to introduce methodological errors in phylogenetic analyses. To mitigate potential artifacts caused by LBA and compositional bias, we re-analyzed the phylogenetic relationships using the amino acid alignment recoded with a reduced amino acid alphabet, the Susko and Roger set of 4 amino acid classes (SR4). This approach diminishes the impact of compositional heterogeneity by grouping amino acids into four classes based on their physicochemical properties (Susko & Roger, 2007) to reduce the impact of multiple substitutions and convergent evolution due to compositional biases. Despite these measures, the placement of Sukunaarchaeum in the ML and BI trees derived from the SR4 recoded dataset remained incongruent. In the SR4-recoded ML tree, Sukunaarchaeum was placed within the Nanobdellati kingdom and weakly supported, specifically as a sister to the clade comprising Aenigmatarchaeota, Nanohalarchaeota, and Huberarchaeota, but the monophyly received only weak UFBP support (UFBP=65, Supplementary Figure S5). This was one of the positions that were not rejected in the AU test. Conversely, the SR4-recoded BI tree suggested a monophyletic relationship between Sukunaarchaeum, Halobacteriota, and Thermoplasmatota, with a low BPP of 0.64 (Supplementary Figure S6). The internal relationships between these three lineages were unresolved (BPP < 0.5), but notably, Sukunaarchaeum was not placed within either of the two phyla.

Finally, to assess the potential impact of rapidly evolving sites within the alignment, we conducted a fast-evolving site removal (FSR) analysis, sequentially removing the fastest-evolving sites from the alignment and re-inferring the phylogeny, on both the original and SR4-recoded alignments. While the FSR analysis did not yield a single consistent result for the branching position of Sukunaarchaeum, it demonstrated relatively stronger support for a monophyletic relationship between Sukunaarchaeum and Nanobdellati (Supplementary Table S5). Notably, the strongest overall support from both SR4 and FSR was for a clade comprising Aenigmatarchaeota, Nanohalarchaeota, Huberarchaeota, and Sukunaarchaeum. However, under specific FSR conditions (30%, 50-70%) and SR4+FSR (40%), Methanobacteriati emerged as a more favored sister group to Sukunaarchaeum than Nanobdellati.

In summary, while a definitive phylogenetic position of Sukunaarchaeum remains elusive, our results robustly support its status as a novel, deeply branching archaeal lineage, distinct from all currently recognized phyla. While a relationship to existing kingdoms, namely Methanobacteriati or Nanobdellati, cannot be entirely excluded, the data strongly suggests that Sukunaarchaeum represents a previously unknown branch of the archaeal tree of life, potentially warranting classification at a high taxonomic rank (e.g., phylum or higher).

### Diversity, Environmental Distribution and Potential Host Association of Sukunaarchaeum Lineage

Beyond its genomic sequence, the biological and ecological characteristics of Sukunaarchaeum, including cell morphology, remain unknown. To gain insights into its distribution and potential ecological role, we analyzed environmental sequence data from marine environments where *C. regius*, the dinoflagellate host from which the Sukunaarchaeum genome was obtained, is found. Utilizing the *Tara* Oceans project, which provides global-scale marine environmental sequence data (de Vargas et al., 2015; Sunagawa et al., 2015), we searched environmental sequence assemblies using the Sukunaarchaeum rRNA gene sequences. While no corresponding sequences were found in the preassembled metagenome data provided by the *Tara* Oceans project or in the OceanDNA MAG catalog (Nishimura & Yoshizawa, 2022), we unexpectedly identified a notable number of homologous sequences exclusively within the *Tara* Oceans metatranscriptome data (The Marine Atlas of *Tara* Oceans Unigenes (MATOU)). While this metatranscriptome assembly was designed to target polyadenylated eukaryotic mRNAs with poly-A tails, archaeal rRNA can be polyadenylated (Evguenieva-Hackenberg et al., 2014), and notably, the Sukunaarchaeum genome encodes proteins involved in rRNA polyadenylation (exosome subunit proteins). Phylogenetic analysis of the rRNA gene sequences obtained from MATOU, alongside sequences representing the overall diversity of archaea, showed that the Sukunaarchaeum-related sequences formed a monophyletic group with the Sukunaarchaeum sequence (Figure 3a; Supplementary Figures S7 and S8), which we designated as the ‘Sukuna-clade’. Notably, the branch lengths within the Sukuna-clade are comparable to those observed within established archaeal kingdoms, such as Nanobdellati and Methanobacteriati, suggesting a similar level of phylogenetic diversity. This suggests that the Sukuna-clade represents a previously overlooked lineage with substantial diversity within the archaeal domain.

**Figure 3.**
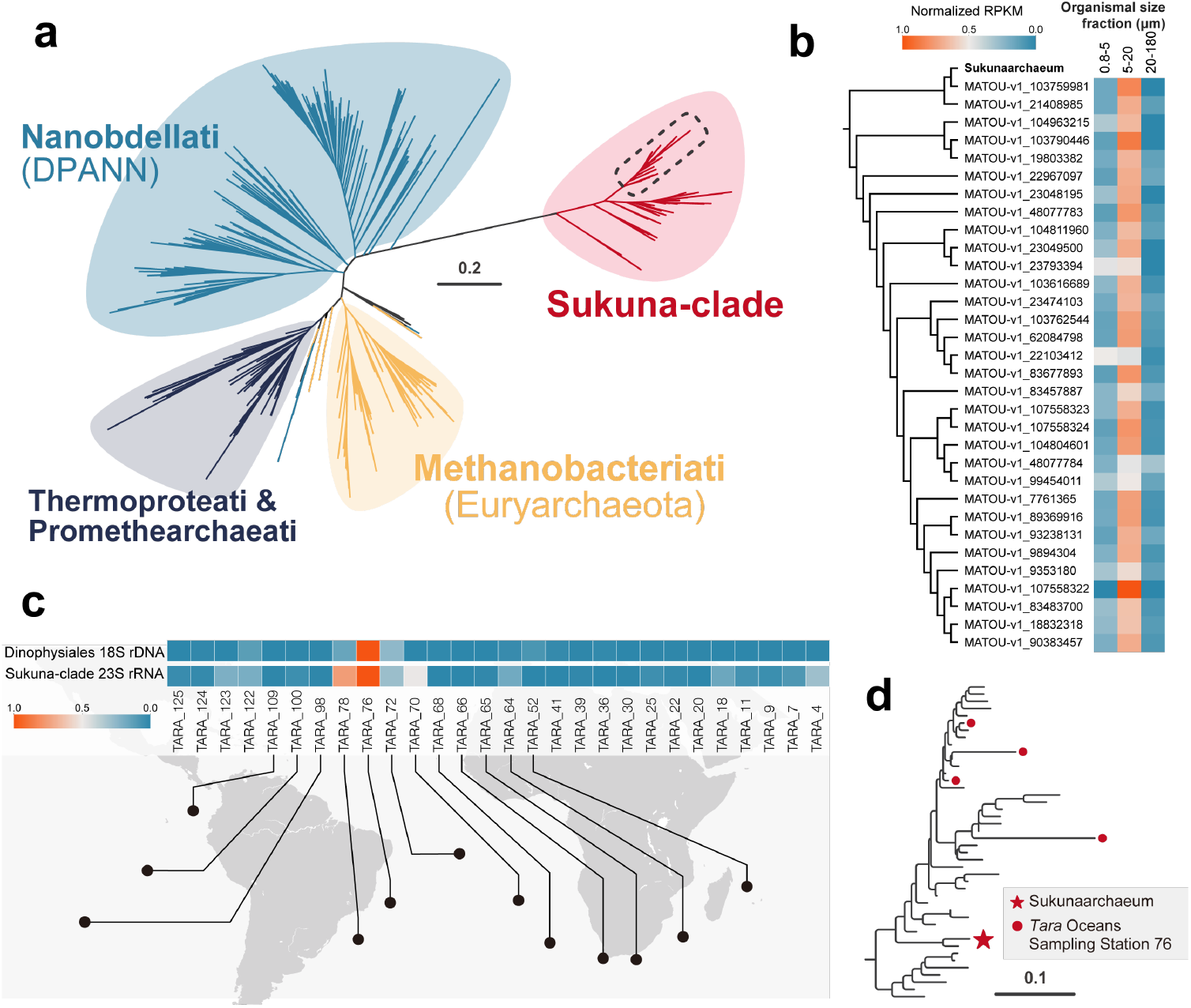
Exploration of Sukunaarchaeum-related sequences in environmental sequence datasets. **a**, Phylogenetic tree of sequences similar to the 23S rRNA gene of Sukunaarchaeum, identified from the *Tara* Oceans metatranscriptome (MATOU) database. **b**, Relative transcript abundance profile of Sukunaarchaeum-related sequences (corresponding to the clade within the dashed line in **a**) in different size fractions from the *Tara* Oceans samples. The cladogram on the left reflects the phylogenetic relationships shown in **a**. The heatmap shows normalized RPKM values, where the sum across the three size fractions (0.8-5 µm, 5-20 µm, and 20-180 µm) equals 1 for each sequence. (See Supplementary Figure S9 for the heatmap including all Sukunaclade sequences shown in **a**). **c**, Relative abundance of Sukunaarchaeum-relatives and Dinophysales across sampling sites. The abundance of Sukunaarchaeum is based on metatranscriptome data, while that of Dinophysales is based on 18S rDNA metabarcoding. **d**, Phylogenetic position of 23S rRNA gene sequences among the ‘Sukuna-clade’ that obtained from *de novo* metagenomic assembly of samples from *Tara* Oceans sampling station 76 (TOSS 76). Scale bars: number of substitutions per site.

We then attempted to gain insights into detailed features of the Sukuna-clade sequences by further utilizing the *Tara* Oceans dataset. Since *Tara* Oceans provides sequence data corresponding to each size range of microorganisms, we first examined the size distribution of Sukunaarchaeum-related sequences. The results revealed that Sukunaarchaeum-related sequences were consistently found more frequently in the larger size fraction (5-20 µm) than in the size fraction (0.8-5 µm) corresponding to the typical cell size of archaea (∼1 µm; Figure 3b; Supplementary Figure S9) (van Wolferen et al., 2022). Next, we examined the frequency of occurrence at each sampling station where the environmental sequences were obtained. We found that these Sukunaarchaeum-related sequences were detected most frequently at a site near the coast of Brazil in the South Atlantic Ocean: *Tara* Oceans sampling station 76 (TOSS 76) (Figure 3c). Consequently, we *de novo* assembled a more comprehensive metagenome assembly by combining multiple metagenomic datasets from TOSS 76. We then searched this assembly for sequences related to the Sukunaarchaeum rRNA gene. As a result, we discovered several partial rRNA gene sequences that formed a monophyletic group with the sequences already obtained from the MATOU, metatranscriptome database (Figure 3d). Read recruitment analysis against these metagenomic sequences further corroborated their prevalence in the 5-20 µm size fraction (Supplementary Table S6). Sequences related to Sukunaarchaeum or its relatives were confirmed in three different types of data: a single-cell genome amplification product, a metatranscriptome, and a metagenome. This result provides strong evidence for the existence of the Sukunaarchaeum lineage.

The abundance of Sukunaarchaeum-related sequences in the 5-20 µm size fraction, typical for many single-celled eukaryotes (Pesant et al., 2015), suggests these archaea rarely exist freely and likely associate closely with larger microbes. This aligns with the discovery of Sukunaarchaeum from the dinoflagellate *C. regius*. Many species within Dinophysales, the order containing *C. regius*, harbor prokaryotic symbionts, mainly comprised of cyanobacteria. Therefore, we analyzed *Tara* Oceans 18S rRNA gene metabarcoding data (de Vargas et al., 2015) and found that Dinophysales sequences were also relatively frequent in the 5-20 µm fraction at TOSS 76, the station where Sukuna-clade sequences peaked (Figure 3c). This co-occurrence might suggest a close symbiotic relationship between Dinophysales and Sukunaarchaeum and its related lineages. However, linking the entire clade primarily to *C. regius* requires careful consideration. Indeed, most described Dinophysales species typically exceed 20 µm in size (Gómez et al., 2011; Hallegraeff et al., 2022; Hernández-Becerril et al., 2008). The cell size of *C. regius* is approximately 40-50 µm at the longest axis of the cell outline (Nakayama et al., 2024), which is larger than the 5-20 µm range, where Sukuna-clade sequences peaked. In fact, previous research has shown that environmental sequences corresponding to the cyanobacteria genome symbiotic with *C. regius* were most frequently found in the 20-180 µm size fraction, with the 5-20 µm fraction being secondary (Nakayama et al., 2024). This size discrepancy highlights the challenge in elucidating the host(s). Considering the significant phylogenetic diversity observed within the Sukuna-clade from environmental surveys, it is highly plausible that different lineages within this clade associate with a variety of hosts. While *Candidatus* Sukunaarchaeum mirabile was discovered in association with *C. regius*, many of its relatives, representing much of the clade’s diversity, may associate with smaller Dinophysales species or perhaps even with entirely different lineages of eukaryotic microorganisms abundant in the 5-20 µm size fraction. Determining the specific host range utilized by the diverse members of the Sukuna-clade remains a key question for future, observation-based research.

### Extreme Genome Reduction and Functional Specialization in Sukunaarchaeum

The Sukunaarchaeum genome displays an extreme functional bias towards genetic information processing (GIP; KEGG category including transcription, translation, replication, and related processes) (Kanehisa et al., 2023). GIP-related genes constitute 52.4% of all protein-coding genes (Figure 4a left; Supplementary Table S7) and 72.6% of those with functional predictions. Conversely, proteins responsible for metabolism were notably absent (Figure 4a middle; Supplementary Table S7), indicating a striking bias in the function of encoded proteins. Expanding the comparison to bacteria, there are some bacterial endosymbionts found in insects that have smaller genomes than that of Sukunaarchaeum. While such extremely reduced genomes also tend to be enriched for GIP functions, Sukunaarchaeum dedicates an even greater proportion of its coding capacity to GIP genes, and a markedly smaller proportion to metabolic functions, compared to these bacterial examples (Figure 4a left, middle). This distinct functional profile suggests that Sukunaarchaeum may have been exposed to extreme genome reduction pressures comparable to those on the bacterial endosymbionts, but that its gene retention was shaped by different selective constraints. This contrast is even clearer when comparing the breakdown of protein function repertoires with *Candidatus* Carsonella ruddii, which has an even smaller genome (159 kbp). While *Ca*. Carsonella ruddii has a relatively large number of genes, including those for amino acid synthesis, carbohydrate metabolism, and energy metabolism (Figure 4b), it possesses fewer genes for certain GIP functions including aminoacyl-tRNA synthetases and protein quality control enzymes compared to Sukunaarchaeum, which retains a more complete GIP toolkit despite its near-total lack of metabolic pathways (Figure 4b).

**Figure 4.**
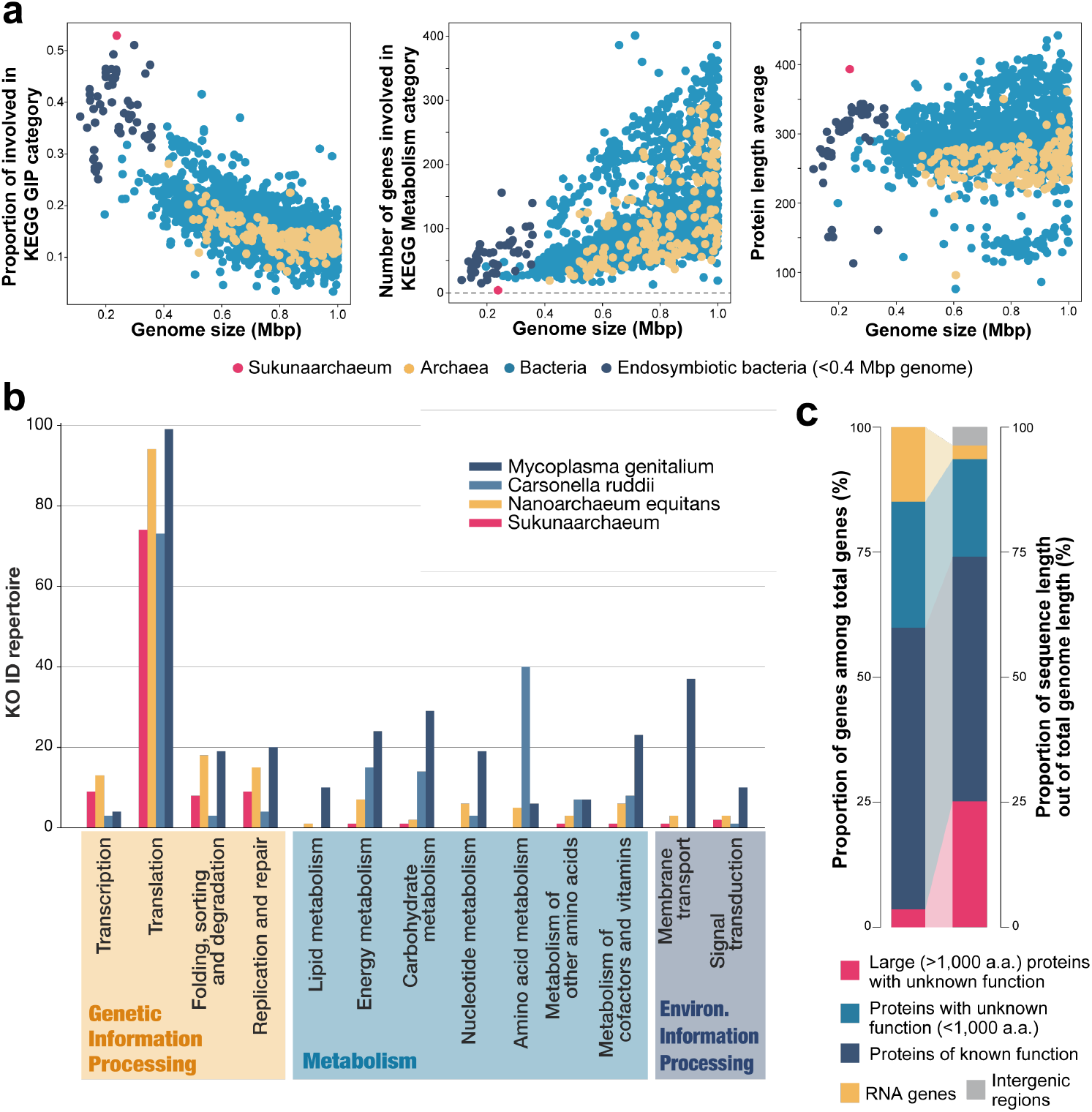
Genomic and proteomic features of Sukunaarchaeum. **a**, Comparative genomic analysis. Scatter plots show genome size versus the proportion of genes in the KEGG Genetic Information Processing (GIP) category (left), the number of genes in the KEGG Metabolism category (middle), and the average protein length (right). **b**, Comparison of gene function repertoires (KEGG categories) across Sukunaarchaeum and selected organisms with small genomes. **c**, Proportion of genes (left) and genomic sequence length (right) attributed to different gene categories in Sukunaarchaeum.

This pattern deviates Sukunaarchaeum genome sharply from known symbiotic genomes. Mutualistic bacterial endosymbionts often lack pathways for independent growth but retain specific biosynthetic capabilities (e.g., essential amino acids, vitamins) that benefit the host (McCutcheon & Moran, 2011; Shigenobu et al., 2000). Even known parasitic bacteria with reduced genomes (e.g., *Mycoplasma, Chlamydia, Rickettsia*) retain more metabolic capacity, including specific biosynthetic and energy-generating pathways, despite their host dependence (Figure 4b) (Andersson et al., 1998; Fraser et al., 1995; Read et al., 2000). The near-complete absence of recognizable metabolic pathways in Sukunaarchaeum precludes any obvious metabolic contribution to its host and strongly suggests an obligate, potentially parasitic, dependence in which it unilaterally exploits host resources. ‘*Nanoarchaeum equitans*’, an archaeon with an extremely small genome of 490 kbp, shows a similar bias towards GIP over metabolism (Figure 4b). However, ‘*N. equitans*’ retains slightly more metabolic capacity than Sukunaarchaeum and the precise relationship with its archaeal host *Ignicoccus hospitalis* remains debated, lacking clear evidence of either parasitism or mutualism (Moissl-Eichinger & Huber, 2011). Sukunaarchaeum’s more profound metabolic depletion compared to ‘*N. equitans*’ suggests an even more extreme level of host dependence, potentially placing a greater burden on its host.

This extreme functional specialization is underscored by the retention of most core protein components essential for transcription, translation, and DNA replication (orthologous to those found across diverse archaea) despite possessing only 189 protein-coding genes (Supplementary Figure S10). Its functionally annotated proteome is thus overwhelmingly dedicated to maintaining and propagating its genetic information. Only five proteins appear clearly unrelated to these core processes: four are transporter subunits (three ABC transporter, one Sec transporter), presumably involved in acquiring essential molecules from the host environment. The fifth shows weak homology to phosphoglycolate phosphatase (PGP; E-value 4e-07, 28.7% identity); however, given the absence of canonical pathways producing its substrate (2-phosphoglycolate) and the low sequence similarity, its true function remains enigmatic and may involve a different substrate entirely. This almost complete loss of metabolic capacity, while retaining the core machinery, represents an unprecedented level of host dependence that distinguishes Sukunaarchaeum even from highly reduced endosymbiotic bacteria (which often retain specific biosynthetic pathways) and the metabolically limited ‘*N. equitans*’.

### Unusually Large Proteins in Sukunaarchaeum: Potential Role in Host Interaction

Despite its highly reduced genome, Sukunaarchaeum encodes several unusually large proteins, a feature uncommon in organisms with such small genomes. The average length of the proteins encoded by the Sukunaarchaeum genome are significantly larger than those encoded by the other genomes with comparable sizes (Figure 4a right; Supplementary Table S7). This characteristic is attributed to a small number of exceptionally large proteins. The largest protein encoded by the genomes of bacteria smaller than 0.4 Mbp is 1,507 amino acids (a.a.), and, among all proteins encoded by the genomes of these symbiotic bacteria with extremely small genomes we analyzed, only 1.5% (197 out of 12,852) exceed 1,000 a.a.. In contrast, 6.9% (13 out of 189) of the proteins encoded by the Sukunaarchaeum genome exceed 1,000 a.a., with the largest reaching 4,756 a.a.. Considering that the median length of Sukunaarchaeum proteins is 224 a.a., these proteins exceeding 1,000 a.a. are exceptionally large. The retention of these large proteins, despite the strong evolutionary pressure for genome reduction expected in Sukunaarchaeum, suggests their crucial roles in the lifestyle of Sukunaarchaeum.

Among the 13 Sukunaarchaeum proteins exceeding 1,000 a.a., homology searches revealed the functions of the five smallest proteins: RNA polymerase subunits, DNA polymerase, topoisomerase, and LonB protease. However, the remaining eight larger proteins showed no apparent sequence similarity to any known proteins, hindering functional inference. Notably, the coding regions for these eight large proteins comprise 25% of the Sukunaarchaeum genome (59,946 out of 238,034 bp; Figure 4c). These large proteins are predicted to be rich in alpha helices, some of which are predicted to be transmembrane helices. TMHMM2.0 (Krogh et al., 2001) and Phobius1.01 (Käll et al., 2004) predicted at least four transmembrane helices for all eight proteins, and DeepTMHMM1.0 (Hallgren et al., 2022) also predicted transmembrane helices for seven proteins. Furthermore, DeepLocPro (Moreno et al., 2024) analysis, which predicts subcellular localization, suggested that all eight large proteins function at the cell membrane (Supplementary Table S2). These results strongly suggest that the large proteins of unknown function in Sukunaarchaeum are localized and function at the cell membrane. Protein structure prediction using AlphaFold2 and 3 did not provide definitive functional insights into the eight large proteins. However, mapping the transmembrane (TM) helix regions predicted by several software onto the predicted three-dimensional structure of one of the large proteins (SKNCREG01_0060) revealed that the TM regions appear to align on a single plane (Supplementary Figure S11). Interestingly, the genes encoding four of the large proteins, including SKNCREG01_0060 (SKN-CREG01_0050-0080), are not only located on the same strand within the genome but are also adjacent to each other, suggesting a functional relationship. These results further support our hypothesis that the large proteins are cellular membrane proteins.

While no homology was identified for the large proteins of Sukunaarchaeum, several parasitic or ectosymbiotic prokaryotes are known to possess large proteins within their reduced genomes. For example, some Nanohalarchaeota species belonging to the cluster 2 of Nanobdellati (DPANN), despite having compact genomes of about 1 Mbp, possess large proteins called SPEAREs, exceeding 5,000 a.a. in size. These are surface proteins thought to play a crucial role in the host invasion process (Hamm et al., 2019; Reva et al., 2023). Additionally, *Candidatus* Nanohalarchaeum antarcticus, a Nanohalarchaeota species with this protein, appears to have a parasitic lifestyle characterized by host cell invasion and lysis (Hamm et al., 2024). Similarly, in bacteria with reduced genomes, there are examples of large proteins localized on the cell surface being involved in host parasitism (Kizina et al., 2022; Seymour et al., 2023).

Given Sukunaarchaeum’s predicted metabolic dependence and the precedent of large, surface-localized proteins mediating host interactions in other prokaryotes, we hypothesize that these eight large proteins are involved in host association. Beyond these eight exceptionally large proteins, many other uncharacterized Sukunaarchaeum proteins are predicted to be membrane-associated. Of the 57 proteins with unknown function, 25 are predicted membrane proteins with an average length of 1,070 a.a., while the remaining 32 are predicted non-membrane proteins with an average length of only 140 a.a. (Supplementary Figure S12). These hypothetical membrane proteins including large proteins may provide clues to the lifestyle of Sukunaarchaeum.

### Sukunaarchaeum: A Minimal Archaeon Genome Challenging Concepts of Cellular Life

The discovery of Sukunaarchaeum not only expands the known boundaries of archaeal diversity but also challenges fundamental concepts of cellular life. The extreme metabolic simplification raises fundamental questions about the minimal requirements for cellular life. Sukunaarchaeum, focused almost entirely on genetic self-perpetuation, represents a compelling example of how far metabolic reduction can proceed within a cellular framework. Its minimal genome, absolute host dependence necessitated by profound metabolic reduction, rapid evolution, and significant investment in large, membrane-associated proteins potentially mediating host interaction constitute a unique combination of characteristics that are collectively reminiscent of viruses. Nonetheless, Sukunaarchaeum remains fundamentally cellular – a key distinction from viruses, which typically lack their own core replication machinery genes and rely on host systems. It possesses ribosomes and the core transcriptional and translational apparatus inherited from cellular ancestors. Thus, while clearly cellular, its extreme metabolic dependence and specialization for self-replication are virus-like in nature, suggesting that Sukunaarchaeum may represent the closest cellular entity discovered to date that approaches a viral strategy of existence.

This extreme evolutionary trajectory challenges existing paradigms of cellular biology. The presence of genes encoding proteins with no detectable homology to known proteins, including the exceptionally large membrane proteins, further highlights the limits of our knowledge and the potential for novel biological functions. Considering the high diversity of Sukunaarchaeum-related lineages revealed through our environmental sequence analysis, further investigation of microbial interactions will likely uncover even more unconventional life forms. The discovery of Sukunaarchaeum illuminates the vast, unexplored biodiversity in microbial interactions, prompting a reevaluation of the minimal functional requirements for cellular organisms and opening new avenues for understanding cellular evolution.

## Materials and Methods

### Sample Collection and DNA Amplification

The Sukunaarchaeum genome analyzed in this study was discovered within a single-cell amplified genome assembly of *Citharistes regius* (cell ID: M16), previously reported by Nakayama et al. (2024) (Nakayama et al., 2024). The *C. regius* M16 cell was collected off the coast of Shimoda, Shizuoka Prefecture, Japan (34°29.222’ N, 139°06.200’ E), using a plankton net with a 25 µm mesh size. The cell was isolated under a microscope using a microcapillary, washed three times with filtered (0.7 µm pore size) and autoclaved seawater, and finally washed once with PCR-grade sterilized water. Whole genome amplification was performed using the REPLI-g Single Cell Kit (QIAGEN) according to the manufacturer’s instructions. The amplified DNA was treated with S1 nuclease (TaKaRa) to digest singlestranded DNA present in the amplified product.

### Library Preparation and Sequencing

For Illumina sequencing, a library was prepared from the amplified and S1 nuclease-treated DNA using the Nextera TruSeq DNA PCR-Free Low Throughput Library Prep Kit (Illumina). The library was sequenced on an Illumina NovaSeq 6000 platform, generating 35.5 million 150 bp paired-end reads. These reads were quality-filtered using fastp (version 0.12.4) with default options, resulting in 33.7 million paired-end reads and a total of 5.1 Gbp. For Oxford Nanopore long-read sequencing, a library was prepared from the amplified DNA using the Rapid Sequencing Kit (SQK-RBK004, Oxford Nanopore Technologies). The library was sequenced on a MinION Flow Cell (R9.4.1) using the MinION platform (Oxford Nanopore Technologies). This yielded reads with a total length of 162 Mb and an N50 of 4.5 kb. Only reads longer than 1 kb (totaling 147 Mb) were used for subsequent assembly.

### Hybrid Genome Assembly

Quality-filtered Illumina short reads and Oxford Nanopore long reads were used for hybrid genome assembly with Unicycler (version 0.4.8) (Wick et al., 2017). A hybrid assembly approach was chosen to leverage the accuracy of short reads and the contiguity of long reads. A preliminary annotation of the entire assembly was performed using the DFAST web service (version 1.6.0) (Tanizawa et al., 2018). Homology searches using rRNA genes detected by DFAST suggested that a 238 kbp circular contig (Contig ID: 5) was likely of archaeal origin, and this contig was selected for further analysis.

### Search for Additional Archaeal Contigs

To search for additional archaeal sequences within the hybrid assembly, we first removed contigs corresponding to known genomes and plasmids: the cyanobacterial symbiont CregCyn (Contigs 1, 2, 6, and 7) (Nakayama et al., 2024), the putative gammaproteobacterial endosymbionts RS3 and XS4 (Contigs 3 and 4) (Nakayama et al., 2025), and the Sukunaarchaeum genome (Contig 5). The remaining 216 contigs (totaling 61 kb) were searched against the NCBI nt database (Sayers et al., 2024) (accessed March 12, 2025) using the blastn web service (Camacho et al., 2009), with an e-value cutoff of 1E-4. In parallel, potential protein-coding sequences were predicted from these remaining contigs using TransDecoder (version 5.7.1; https://github.com/TransDecoder/TransDecoder). All open reading frames (ORFs) encoding at least 50 amino acids were predicted using the default settings (no strand-specific option). The resulting 285 predicted protein sequences were searched against the ClusteredNR database (nr_clustered, accessed March 12, 2025) using the blastp web service, with an e-value cutoff of 1E-4. The results of the blastn and blastp searches are summarized in Table S3. No sequences other than the already identified Sukunaarchaeum genome showed significant homology to archaeal sequences.

### Comparative Genome Assembly Analysis

To evaluate the robustness and consistency of the Sukunaarchaeum genome assembly, multiple independent assemblies were performed and compared using quality-filtered Illumina paired-end reads and Oxford Nanopore long reads obtained from the single-cell amplified genome product.

Short-read assembly: Subsets representing 10%, 50%, and 100% of the total Illumina reads were randomly sampled using the sample command of seqkit (version 2.2.0) (Shen et al., 2024). Assemblies were performed on these subsets using three different assemblers: MEGAHIT (version 1.2.9) (Li et al., 2015), metaSPAdes (version 3.15.5) (Nurk et al., 2017; Prjibelski et al., 2020), and Unicycler (version 0.5.1) (Wick et al., 2017).

Long-read preprocessing and assembly: Nanopore reads were prepared in two ways. First, reads were quality-filtered and trimmed using Filtlong (version 0.2.1; https://github.com/rrwick/Filtlong), referencing Illumina reads (options: ‘--trim --split 10 -- min_length 1000 --mean_q_weight 10’), to create ‘uncorrected’ long reads. Second, raw Nanopore reads were error-corrected using Illumina reads with Ratatosk (version 0.9.0) (Holley et al., 2021), and the corrected reads were subsequently processed with Filtlong using the same parameters to create ‘corrected’ long reads. Assemblies were performed on both the ‘uncorrected’ and ‘corrected’ long read sets using three assemblers: Flye (version 2.9) (Kolmogorov et al., 2019), Raven (version 1.8.3) (Vaser & Šikić, 2021), and metaMDBG (version 1.0) (Benoit et al., 2024).

Contigs corresponding to the Sukunaarchaeum genome were identified and extracted from each assembly result. Specifically, for each assembly result, blastn (version 2.12.0) was executed using the Sukunaarchaeum genome sequence (derived from the hybrid assembly used for the main analysis in this study; referred to here as the reference) as the query. Contigs showing high similarity (identity ≥ 98% and bitscore ≥ 5000) were extracted as Sukunaarchaeum genome contigs. Each extracted Sukunaarchaeum contig, along with the reference contig from the hybrid assembly, was evaluated using QUAST (version 5.3.0) (Mikheenko et al., 2023), and comparative assembly statistics (Supplementary Table S1) were calculated.

### Genome Annotation

Structural annotation of the 238 kbp genome was performed by manual curation, integrating results from DFAST (version 1.2.16) and TransDecoder longorfs (version 5.5.0) results. Preliminary protein functional annotation was performed via sequence similarity searches using PSI-BLAST with e-value cutoffs of 1E-4 against the GenBank nr database (accessed at June 1, 2023) and domain prediction using InterProScan (version 5.55-88.0) (Jones et al., 2014). In addition, the tertiary structures of all proteins were predicted by AlphaFold (versions 2.2.0, 2.3.2, 3) (Abramson et al., 2024; Jumper et al., 2021). The resulting structural models were used in structural similarity searches using the Foldseek web server (van Kempen et al., 2024) (https://search.foldseek.com/ search). Eleven proteins were functionally annotated based on structural similarity, despite lacking detectable primary sequence similarity. KEGG Orthology (KO) IDs were assigned to each protein based on the integrated functional annotation, incorporating results from GhostKOALA and BlastKOALA (Kanehisa et al., 2016).

### CheckM2 Analysis

CheckM2 (Chklovski et al., 2023) quality assessment was adapted for the Sukunaarchaeum genome. Typically, CheckM2 annotates protein-coding genes with KO IDs by similarity search against UniRef100 and converts these annotations to a feature vector. As most of the protein encoded by Sukunaarchaeum genome can not be linked to KO IDs by the default method, we prepared an annotation file containing our curated KO ID assignments in the CheckM2 intermediate format. A modified ‘predictQuality.py’ script (provided by CheckM2) was used to read this custom annotation file, convert a feature vector, and calculate genome completeness and contamination. The assigned KO IDs were subjected to the CheckM2 (version 1.0.2) analysis, resulting in 83.68% completeness and 0.18% contamination.

To further assess the validity of this completeness, a comparative analysis was performed: Archaeal genomes of smaller than 1 Mb consisting a single contig were selected from Genome Taxonomy Database (GTDB) r220 (Parks et al., 2022). Genome fragments of 240 kb were generated through a sliding window approach with window size = 240 kb and slide width = 10 kb. The resultant genome fragments were submitted to CheckM2 and evaluated their completeness and contamination.

### Genome Dataset Collection and Processing

For comparative genomic analyses, all 6519 prokaryotic genomes smaller than 1 Mbp were downloaded from GenBank on December 15, 2023. Of these, annotations were available for 4574 genomes. The remaining 1945 genomes lacking annotations were newly annotated using DFAST (version 1.2.16). To exclude genomes with low completeness, likely representing partial genome assemblies, 2828 genomes with at least two rRNAs and 18 tRNAs were retained for further analysis. For functional comparison, all proteins in these retained genomes were assigned KO IDs by GhostKOALA. For the 87 genomes smaller than 0.4 Mbp, we manually checked their BioSample information in NCBI to determine if they were reported as endosymbionts. We found that 57 were genomes of endosymbionts while the others were metagenome-assembled genomes (MAGs). Genomic statistics for this dataset, as used for the scatterplots are available in Supplementary Table S7.

### Phylogenetic Analysis of 70 genes

To elucidate the phylogenetic position of Sukunaarchaeum, we prepared the multigene dataset based on the 151 archaeal marker genes used in a previous study (Dombrowski et al., 2020). For each of the 151 genes, we selected the homologous sequences by blastp searches from the predicted proteome of Sukunaarchaeum. The 70 genes for which homologous sequences were detected from Sukunaarchaeum were selected for the subsequent phylogenetic analysis. To sample taxa reflecting the diversity of Archaea, we selected a single genome with the highest completeness from each archaeal order leaving 149 genomes for alignment. In this analysis, we adopted the archaeal taxonomy of the GTDB r214. The proteome of each genome was predicted using DFAST (version 1.2.16) and homologous sequences of the 70 marker genes were retrieved by blastp. We aligned each of the 70 protein sequences using MAFFT (version 7.490) (Katoh & Standley, 2013) with the L-INS-i method and trimmed ambiguously aligned positions with BMGE (version 2.0) (Criscuolo & Gribaldo, 2010). The remaining sites of the 70 proteins were concatenated into a single alignment comprising 150 taxa and 18,286 amino acid positions. We inferred the maximum likelihood (ML) tree based on the alignment by using IQ-TREE (version 2.2.2.5) (Minh et al., 2020) under the LG+C60+F+I+R10 model, which was selected by adding C60 to the results of a model test performed on homogeneous models using ‘-mset LG,WAG,Q.pfam’ option. The statistical support for each bipartition in the ML tree was calculated by 1000-replicate ultrafast bootstrap approximation (Hoang et al., 2018) and 100-replicate nonparametric bootstrap analysis with the ML tree used as the guide for estimating PMSF parameters (Wang et al., 2018). The alignment was also subjected to Bayesian analysis with the GTR+CAT+Γ model using PhyloBayes (version 1.9) (Lartillot et al., 2013). We ran two Markov chain Monte Carlo runs for 30,000 cycles with a burn-in of the first 3,000 cycles. Although the two chains did not converge, inspection revealed no difference in the inferred phylogenetic position of Sukunaarchaeum between the chains.

We evaluated the possibility of Sukunaarchaeum branching at alternative positions using the approximately unbiased (AU) test (Shimodaira, 2002). We generated a set of the alternative trees by pruning the Sukunaarchaeum branch from the ML tree and regrafting it onto each possible internal and terminal branch of the remaining tree. A total of 295 trees, including the ML tree, were subjected to the AU test using IQ-TREE with the LG+C60+F+I+R10 model. As a result, 16 trees, including the ML tree, were not rejected at the 5% significance level, providing candidate phylogenetic positions from which Sukunaarchaeum could branch.

To reduce compositional biases and the effect of long branch attraction, we recoded the alignment into four characters using the Susko and Roger set of 4 amino acid classes (SR4) (Susko & Roger, 2007). ML tree was inferred under the GTR+SR4C60+F+I+R8 model, where SR4C60 represents the profiles of SR4 recoded frequencies created by adding up the 20 amino acid frequencies of the original C60 profiles. The statistical support for each bipartition in the ML tree was calculated by 1000-replicate ultrafast bootstrap approximation. Bayesian inference was also performed on the SR4 recoded alignment in the same manner as for the original alignment. Two chains did not converge, but yielded consistent placement for Sukunaarchaeum.

We evaluated the contribution of fast-evolving sites in the alignments to the phylogenetic inference by performing fast-evolving sites removal (FSR) analyses. The substitution rate of each site was estimated across the original alignment and ML tree using IQ-TREE2 with the ‘-wsr’ option. For both the original alignment and the SR4 recoded alignment, eight sub-alignments each were generated by stepwise removal of the fastest evolving sites by 10% increments. For each alignment, the ML tree was reconstructed after model testing (Supplementary Table S8) and the support for the phylogenetic position of Sukunaarchaeum was assessed by 1000-replicate ultrafast bootstrap trees.

### Search for Sukunaarchaeum-Related rRNA Sequences from Metatranscriptomic Data

To identify Sukunaarchaeum-related rRNA gene sequences, we performed BLASTN searches against several publicly available sequence datasets, including both metagenomic and metatranscriptomic assemblies:

- *Tara* Oceans Eukaryotic Gene Catalog (MATOU-v1): Metatranscriptomic unigene dataset (Carradec et al., 2018). MATOU-v1.fna.gz file was downloaded from the archived Genoscope *Tara* Oceans website (https://www.genoscope.cns.fr/tara/localdata/data/Geneset-v1/).
- Ocean Microbial Reference Gene Catalog v2 (OM-RGC.v2): Catalog of genes from marine microbes generated by the *Tara* Oceans project (Salazar et al., 2019). OM-RGC_v2.tsv.gz file was obtained from the EBI BioStudies database (accession S-BSST297; https://www.ebi.ac.uk/biostudies/studies/S-BSST297). All the assemblies were used for the search.
- *Tara* Oceans Binned Genome Dataset : Collection of 2,631 draft microbial genomes assembled from *Tara* Oceans global metagenomes (Tully et al., 2018). A compressed file of the dataset was downloaded from https://figshare.com/ndownloader/articles/5188273/versions/4. For the Sukunaarchaeum-related rRNA sequence search, all of the primary contigs in the dataset were used.
- *Tara* Oceans Eukaryotic Genomes (SMAGs): Curated *Tara* Oceans single-cell and metagenome assembled genomes (Delmont et al., 2022). Assemblies_1000nt.tar.gz file was obtained from the Genoscope *Tara* Oceans website (https://www.genoscope.cns.fr/tara/localdata/data/SMAGs-v1/).
- The OceanDNA MAG catalog: Metagenome catalog of over 50,000 prokaryotes reconstructed from various marine environments (Nishimura & Yoshizawa, 2022). Both the representative and non-representative MAGs were downloaded from figshare (https://doi.org/10.6084/m9.figshare.c.5564844.v1).

The blastn (version 2.12.0) searches against each database were carried out using the Sukunaarchaeum rRNA gene sequences (16S and 23S rRNA gene) as the query, with an e-value cutoff of 1E-50. Only the MATOU-v1 dataset returned sequences with significant similarity to the query.

### Expression Pattern of Sukunaarchaeum rRNA Gene in Metatranscriptome Data

To assess the expression (as a proxy for abundance and activity) of Sukunaarchaeum-related rRNA genes across different size fractions and geographic locations, we analyzed expression data for sequences in MATOU-v1 (Carradec et al., 2018). RPKM (Reads Per Kilobase Million) values for each Sukuna-clade sequence in each sample were obtained from the metatranscriptomic_occurrences.tsv.gz file, available at https://www.genoscope.fr/tara/localdata/data/Geneset-v1/. *Tara* Oceans sample metadata were obtained from Carradec *et al*. (2018) (Carradec et al., 2018), Supplementary Data 8.

The *Tara* Oceans project collected metatranscriptomic data from multiple size fractions. We focused on three contiguous filter size fractions: 0.8-5 µm, 5-20 µm, and 20-180 µm, selected for their continuous size ranges and the relatively high availability of metatranscriptomic data across sampling stations. To ensure robust comparisons and minimize biases due to uneven sampling, we identified *Tara* Oceans sampling stations with expression data available for all three selected filter sizes. For each of these stations, we summed the RPKM values for each of the three size fractions and calculated the mean RPKM values. This aggregation was performed separately for samples from two depth categories, the surface (SRF) and the deep chlorophyll maximum (DCM). RPKM values for the three size fractions (0.8-5 µm, 5-20 µm, and 20-180 µm) were available for the SRF data of 19 stations and the DCM data of 7 stations. The SRF and DCM data were then integrated by calculating the mean value for overlapping sampling stations, yielding a representative RPKM value for each sequence in each size fraction. To determine the relative expression of each sequence across the three size fractions, we normalized the RPKM values such that the sum of the expression levels across the three fractions for each sequence equaled 1. This normalization allows for comparison of relative abundance across size fractions, accounting for differences in sequencing depth.

To determine which sampling stations showed the highest expression of Sukuna-clade sequences, we focused on the 5-20 µm size fraction, which exhibited relatively high expression levels of Sukuna-clade rRNA. For each sampling station, we calculated the mean RPKM value for each sequence in both the SRF and DCM depths. Finally, we calculated the mean RPKM value across all Sukuna-clade sequences for each station.

### Occurrence Pattern of Dinophysales Dinoflagellates Across Sampling Stations

To examine the relative abundance of Dinophysales dinoflagellates, a group known to harbor prokaryotic symbionts and the order to which the host *Citharistes regius* belongs, we analyzed the *Tara* Oceans 18S rRNA gene V9 metabarcode data (de Vargas et al., 2015). We used the Database_W4_barcode_occurences.tsv file, which is distributed at https://doi.pangaea.de/10.1594/ PANGAEA.843018. This TSV file contains taxonomic information and read counts for each barcode sequence in each metabarcoding sample from the *Tara* Oceans V9 dataset. From this file, we extracted information for barcode sequences assigned to the order Dinophysales (using the taxonomic assignment “Dinophysiales” provided in the dataset, noting that this is a taxonomic synonym). Separately, for all barcode sequences listed in the Database_W4_barcode_occurences.tsv file, we summed the read counts in each metabarcoding sample to obtain the total number of barcode sequences in each sample.

For each metabarcoding sample, we calculated the relative abundance of Dinophysales sequences by dividing the total read count of Dinophysales sequences by the total read count of all barcode sequences in that sample. Then, for each sampling station, we calculated the average relative abundance of Dinophysales sequences. To facilitate comparison with the expression data of the Sukuna-clade 23S rRNA gene, we focused on the 5-20 µm size fraction and calculated the average relative abundance of Dinophysales sequences at each station within this size fraction.

### Metagenome Assembling and Search for Sukunaarchaeum-related sequences

To investigate the presence and genomic context of Sukunaarchaeum-related lineages at the site of their highest transcriptomic signal (*Tara* Oceans sampling station 76, TOSS 76), we performed a targeted *de novo* metagenome co-assembly. Raw Illumina sequence reads were downloaded from the NCBI Short Read Archive (SRA) for 12 datasets corresponding to TOSS 76 size fractions (0.8-5 µm, 5-20 µm, and 20-180 µm; SRA accessions provided in Supplementary Table S9). Reads were quality-filtered using fastp (version 0.23.2) (Chen et al., 2018) with the default option. All filtered reads were then co-assembled using MEGAHIT (version 1.2.9) (Li et al., 2015) with the default option. We searched the resulting TOSS 76 metagenomic assembly for Sukunaarchaeum-related sequences using blastn (version 2.15.0) with the 16S and 23S rRNA gene sequences from Sukunaarchaeum as queries. This search identified five potential Sukunaarchaeum-related contigs: one 399 bp contig showing similarity to the 16S rRNA gene and four contigs (313-642 bp) similar to the 23S rRNA gene (E-values < 1E-50).

To assess the abundance and origin of reads corresponding to these contigs within the different size-fraction datasets, we conducted a read recruitment analysis. Read1 files from each metagenomic dataset (total read counts used are listed in Supplementary Table S9) were used as queries in a blastn (version 2.12.0) search against the five identified rRNA contigs. Reads yielding hits with ≥ 99% identity and an alignment length of ≥ 100 bp were counted as recruited reads corresponding to that contig. Recruited reads corresponding to these contigs were detected predominantly in datasets from the 5-20 µm size fraction (four datasets) and one dataset from the 20-180 µm size fraction, while reads from the 0.8-5 µm datasets did not meet these recruitment criteria (see Supplementary Table S6 for detailed recruitment counts per contig and dataset).

### Phylogenetic Analysis rRNA

To elucidate the phylogenetic relationship between Sukunaarchaeum and its related sequences detected from *Tara* Oceans data, we conducted phylogenetic analyses using 16S and 23S rRNA sequences. To generate a reference dataset, we selected a single genome with the highest completeness from each archaeal family from the GTDB r214, leaving 396 genomes. 16S and 23S rRNA genes of each genome were predicted using DFAST (version 1.2.16) and curated manually, resulting in 294 unique 16S rRNA gene sequences and 279 unique 23S rRNA gene sequences. We aligned the 16S and 23S sequences, respectively, along with the homologous sequences from Sukunaarchaeum and the related sequences recovered from the MATOU dataset, using MAFFT (version 7.490) with the L-INS-i method and the ‘--adjustdirectionaccurately’ option. Ambiguously aligned sites were trimmed using BMGE (version 2.0) with the default option. The alignments comprised 306 taxa and 1,387 nucleotide positions (16S rRNA) and 371 taxa and 2,540 nucleotide positions (23S rRNA). ML trees were inferred from each alignment using IQ-TREE (version 2.2.2.5) under the GTR+F+G model with 1000-replicate ultrafast bootstrap approximation. Subsequently, we added the partial rRNA gene sequences (one 16S, four 23S) obtained from our TOSS 76 *de novo* metagenomic assembly to the respective datasets and repeated the same analysis. The alignments including TOSS 76 sequences comprised 307 taxa and 1,421 nucleotide positions (16S rRNA) and 375 taxa and 2663 nucleotide positions (23S rRNA).

## Acknowledgment

This work was supported by JSPS KAKENHI Grant Numbers JP16H06280, 20H03305, 18KK0203, 21K15131, 22KJ0401, 23K27226, 24K09587 and BPI06050, Institute for Fermentation, Osaka (IFO) (Grant ID: G-2025-2-067), as well as by the World Premier International Research Center Initiative (WPI Initiative), MEXT, Japan. R.H. was supported by JSPS Overseas Research Fellowships. We thank the *Tara* Oceans consortium and sponsors who supported the *Tara* Oceans Expedition for making the data accessible. Computations in this study were partially performed on the NIG supercomputer at ROIS National Institute of Genetics. The manuscript file uploaded to bioR*χ*iv was generated using the LaTeX template adapted by Stephen Royle (https://github.com/quantixed) available at https://github.com/quantixed/manuscripttemplates.

## Data Availability

The annotated genome sequence of *Ca*. Sukunaarchaeum mirabile has been deposited in DDBJ/ENA/GenBank under accession number AP040136. Raw reads of single-cell amplified genomes of a *Citharistes regius* cell are available on the Sequence Read Archives (DRA/SRA/ENA: accession numbers DRR628309 and DRR628310). Supporting data including files for phylogenetic analyses, genome assembly, and protein structure models are available on Zenodo (DOI: 10.5281/zenodo.15271943).

## References

Abramson, J., Adler, J., Dunger, J., Evans, R., Green, T., Pritzel, A., Ronneberger, O., Willmore, L., Ballard, A. J., Bambrick, J., Bodenstein, S. W., Evans, D. A., Hung, C.-C., O’Neill, M., Reiman, D., Tunyasuvunakool, K., Wu, Z., Žemgulytė, A., Arvaniti, E., … Jumper, J. M. (2024). Accurate structure prediction of biomolecular interactions with AlphaFold 3. Nature, 630(8016), 493–500.

Andersson, S. G., Zomorodipour, A., Andersson, J. O., Sicheritz-Pontén, T., Alsmark, U. C., Podowski, R. M., Näslund, A. K., Eriksson, A. S., Winkler, H. H., & Kurland, C. G. (1998). The genome sequence of Rickettsia prowazekii and the origin of mitochondria. Nature, 396(6707), 133–140.

Baños, H., Susko, E., & Roger, A. J. (2024). Is over-parameterization a problem for profile mixture models? Systematic Biology, 73(1), 53–75.

Bar-On, Y. M., & Milo, R. (2019). The biomass composition of the oceans: A blueprint of our blue planet. Cell, 179(7), 1451–1454.

Benoit, G., Raguideau, S., James, R., Phillippy, A. M., Chikhi, R., & Quince, C. (2024). High-quality metagenome assembly from long accurate reads with metaMDBG. Nature Biotechnology, 42(9), 1378–1383.

Bowers, R. M., Kyrpides, N. C., Stepanauskas, R., Harmon-Smith, M., Doud, D., Reddy, T. B. K., Schulz, F., Jarett, J., Rivers, A. R., Eloe-Fadrosh, E. A., Tringe, S. G., Ivanova, N. N., Copeland, A., Clum, A., Becraft, E. D., Malmstrom, R. R., Birren, B., Podar, M., Bork, P., … Woyke, T. (2017). Minimum information about a single amplified genome (MISAG) and a metagenome-assembled genome (MIMAG) of bacteria and archaea. Nature Biotechnology, 35(8), 725–731.

Camacho, C., Coulouris, G., Avagyan, V., Ma, N., Papadopoulos, J., Bealer, K., & Madden, T. L. (2009). BLAST+: architecture and applications. BMC Bioinformatics, 10(1), 421.

Carradec, Q., Coordinators, T. O., Pelletier, E., Da Silva, C., Alberti, A., Seeleuthner, Y., Blanc-Mathieu, R., Lima-Mendez, G., Rocha, F., Tirichine, L., Labadie, K., Kirilovsky, A., Bertrand, A., Engelen, S., Madoui, M.-A., Méheust, R., Poulain, J., Romac, S., Richter, D. J., … Wincker, P. (2018). A global ocean atlas of eukaryotic genes. Nature Communications, 9(1). 10.1038/s41467-017-02342-1

Castelle, C. J., Brown, C. T., Anantharaman, K., Probst, A. J., Huang, R. H., & Banfield, J. F. (2018). Biosynthetic capacity, metabolic variety and unusual biology in the CPR and DPANN radiations. Nature Reviews. Microbiology, 16(10), 629–645.

Chen, S., Zhou, Y., Chen, Y., & Gu, J. (2018). fastp: an ultra-fast all-in-one FASTQ preprocessor. Bioinformatics (Oxford, England), 34(17), i884–i890.

Chklovski, A., Parks, D. H., Woodcroft, B. J., & Tyson, G. W. (2023). CheckM2: a rapid, scalable and accurate tool for assessing microbial genome quality using machine learning. Nature Methods, 20(8), 1203–1212.

Coale, T. H., Loconte, V., Turk-Kubo, K. A., Vanslembrouck, B., Mak, W. K. E., Cheung, S., Ekman, A., Chen, J.-H., Hagino, K., Takano, Y., Nishimura, T., Adachi, M., Le Gros, M., Larabell, C., & Zehr, J. P. (2024). Nitrogen-fixing organelle in a marine alga. Science (New York, N.Y.), 384(6692), 217–222.

Criscuolo, A., & Gribaldo, S. (2010). BMGE (Block Mapping and Gathering with Entropy): a new software for selection of phylogenetic informative regions from multiple sequence alignments. BMC Evolutionary Biology, 10, 210.

Cross, K. L., Campbell, J. H., Balachandran, M., Campbell, A. G., Cooper, C. J., Griffen, A., Heaton, M., Joshi, S., Klingeman, D., Leys, E., Yang, Z., Parks, J. M., & Podar, M. (2019). Targeted isolation and cultivation of uncultivated bacteria by reverse genomics. Nature Biotechnology, 37 (11), 1314–1321.

de Vargas, C., Audic, S., Henry, N., Decelle, J., Mahé, F., Logares, R., Lara, E., Berney, C., Le Bescot, N., Probert, I., Carmichael, M., Poulain, J., Romac, S., Colin, S., Aury, J.-M., Bittner, L., Chaffron, S., Dunthorn, M., Engelen, S., … Karsenti, E. (2015). Eukaryotic plankton diversity in the sunlit ocean. Science (New York, N.Y.), 348(6237), 1261605.

Delmont, T. O., Gaia, M., Hinsinger, D. D., Frémont, P., Vanni, C., Fernandez-Guerra, A., Eren, A. M., Kourlaiev, A., d’Agata, L., Clayssen, Q., Villar, E., Labadie, K., Cruaud, C., Poulain, J., Da Silva, C., Wessner, M., Noel, B., Aury, J.-M., Tara Oceans Coordinators, … Jaillon, O. (2022). Functional repertoire convergence of distantly related eukaryotic plankton lineages abundant in the sunlit ocean. Cell Genomics, 2(5), 100123.

Dombrowski, N., Williams, T. A., Sun, J., Woodcroft, B. J., Lee, J.-H., Minh, B. Q., Rinke, C., & Spang, A. (2020). Undinarchaeota illuminate DPANN phylogeny and the impact of gene transfer on archaeal evolution. Nature Communications, 11(1), 3939.

Eme, L., Tamarit, D., Caceres, E. F., Stairs, C. W., De Anda, V., Schön, M. E., Seitz, K. W., Dombrowski, N., Lewis, W. H., Homa, F., Saw, J. H., Lombard, J., Nunoura, T., Li, W.-J., Hua, Z.-S., Chen, L.-X., Banfield, J. F., John, E. S., Reysenbach, A.-L., … Ettema, T. J. G. (2023). Inference and reconstruction of the heimdallarchaeial ancestry of eukaryotes. Nature, 618(7967), 992–999.

Evguenieva-Hackenberg, E., Hou, L., Glaeser, S., & Klug, G. (2014). Structure and function of the archaeal exosome. Wiley Interdisciplinary Reviews. RNA, 5(5), 623–635.

Felsenstein, J. (1978). Cases in which parsimony or compatibility methods will be positively misleading. Systematic Biology, 27 (4), 401–410.

Fraser, C. M., Gocayne, J. D., White, O., Adams, M. D., Clayton, R. A., Fleischmann, R. D., Bult, C. J., Kerlavage, A. R., Sutton, G., Kelley, J. M., Fritchman, R. D., Weidman, J. F., Small, K. V., Sandusky, M., Fuhrmann, J., Nguyen, D., Utterback, T. R., Saudek, D. M., Phillips, C. A., … Venter, J. C. (1995). The minimal gene complement of Mycoplasma genitalium. Science (New York, N.Y.), 270(5235), 397–403.

Golyshina, O. V., Toshchakov, S. V., Makarova, K. S., Gavrilov, S. N., Korzhenkov, A. A., La Cono, V., Arcadi, E., Nechitaylo, T. Y., Ferrer, M., Kublanov, I. V., Wolf, Y. I., Yakimov, M. M., & Golyshin, P. N. (2017). “ARMAN” archaea depend on association with euryarchaeal host in culture and in situ. Nature Communications, 8(1), 60.

Gómez, F., López-García, P., & Moreira, D. (2011). Molecular phylogeny of dinophysoid dinoflagellates: The systematic position of Oxyphysis oxytoxoides and the Dinophysis hastata group (dinophysales, Dinophyceae)(1): Molecular phylogeny of dinophysales. Journal of Phycology, 47 (2), 393–406.

Hallegraeff, G. M., Eriksen, R. S., Davies, C. H., & Uribe-Palomino, J. (2022). Marine planktonic dinophysoid dinoflagellates (order Dinophysales): 60 years of species-level distributions in Australian waters. Australian Systematic Botany, 35(6), 469–500.

Hallgren, J., Tsirigos, K. D., Pedersen, M. D., Almagro Armenteros, J. J., Marcatili, P., Nielsen, H., Krogh, A., & Winther, O. (2022). DeepTMHMM predicts alpha and beta transmembrane proteins using deep neural networks. In bioRxiv. 10.1101/2022.04.08.487609

Hamm, J. N., Erdmann, S., Eloe-Fadrosh, E. A., Angeloni, A., Zhong, L., Brownlee, C., Williams, T. J., Barton, K., Carswell, S., Smith, M. A., Brazendale, S., Hancock, A. M., Allen, M. A., Raftery, M. J., & Cavicchioli, R. (2019). Unexpected host dependency of Antarctic Nanohaloarchaeota. Proceedings of the National Academy of Sciences of the United States of America, 116(29), 14661–14670.

Hamm, J. N., Liao, Y., von Kügelgen, A., Dombrowski, N., Landers, E., Brownlee, C., Johansson, E. M. V., Whan, R. M., Baker, M. A. B., Baum, B., Bharat, T. A. M., Duggin, I. G., Spang, A., & Cavicchioli, R. (2024). The parasitic lifestyle of an archaeal symbiont. Nature Communications, 15(1), 6449.

He, X., McLean, J. S., Edlund, A., Yooseph, S., Hall, A. P., Liu, S.-Y., Dorrestein, P. C., Esquenazi, E., Hunter, R. C., Cheng, G., Nelson, K. E., Lux, R., & Shi, W. (2015). Cultivation of a human-associated TM7 phylotype reveals a reduced genome and epibiotic parasitic lifestyle. Proceedings of the National Academy of Sciences of the United States of America, 112(1), 244–249.

Hernández-Becerril, D. U., Ceballos-Corona, J. G. A., Esqueda-Lara, K., Tovar-Salazar, M. A., & León-Álvarez, D. (2008). Marine planktonic dinoflagellates of the order Dinophysiales (Dinophyta) from coasts of the tropical Mexican Pacific, including two new species of the genus Amphisolenia. Journal of the Marine Biological Association of the United Kingdom. Marine Biological Association of the United Kingdom, 88(1), 1–15.

Hoang, D. T., Chernomor, O., von Haeseler, A., Minh, B. Q., & Vinh, L. S. (2018). UFBoot2: Improving the ultra-fast bootstrap approximation. Molecular Biology and Evolution, 35(2), 518–522.

Holley, G., Beyter, D., Ingimundardottir, H., Møller, P. L., Kristmundsdottir, S., Eggertsson, H. P., & Halldorsson, B. V. (2021). Ratatosk: hybrid error correction of long reads enables accurate variant calling and assembly. Genome Biology, 22(1), 28.

Huber, H., Hohn, M. J., Rachel, R., Fuchs, T., Wimmer, V. C., & Stetter, K. O. (2002). A new phylum of Archaea represented by a nanosized hyperthermophilic symbiont. Nature, 417 (6884), 63–67.

Hug, L. A., Baker, B. J., Anantharaman, K., Brown, C. T., Probst, A. J., Castelle, C. J., Butterfield, C. N., Hernsdorf, A. W., Amano, Y., Ise, K., Suzuki, Y., Dudek, N., Relman, D. A., Finstad, K. M., Amundson, R., Thomas, B. C., & Banfield, J. F. (2016). A new view of the tree of life. Nature Microbiology, 1, 16048.

Husnik, F., Tashyreva, D., Boscaro, V., George, E. E., Lukeš, J., & Keeling, P. J. (2021). Bacterial and archaeal symbioses with protists. Current Biology: CB, 31(13), R862–R877.

Imachi, H., Nobu, M. K., Ishii, S., Hirakata, Y., Ikuta, T., Isaji, Y., Miyata, M., Miyazaki, M., Morono, Y., Murata, K., Nakagawa, S., Ogawara, M., Okada, S., Saito, Y., Sakai, S., Shimamura, S., Tahara, Y. O., Takaki, Y., Takano, Y., … Takai, K. (2025). Eukaryotes’ closest relatives are internally simple syntrophic archaea. In bioRxiv. 10.1101/2025.02.26.640444

Imachi, H., Nobu, M. K., Nakahara, N., Morono, Y., Ogawara, M., Takaki, Y., Takano, Y., Uematsu, K., Ikuta, T., Ito, M., Matsui, Y., Miyazaki, M., Murata, K., Saito, Y., Sakai, S., Song, C., Tasumi, E., Yamanaka, Y., Yamaguchi, T., … Takai, K. (2020). Isolation of an archaeon at the prokaryote-eukaryote interface. Nature, 577 (7791), 519–525.

Jones, P., Binns, D., Chang, H.-Y., Fraser, M., Li, W., McAnulla, C., McWilliam, H., Maslen, J., Mitchell, A., Nuka, G., Pesseat, S., Quinn, A. F., Sangrador-Vegas, A., Scheremetjew, M., Yong, S.-Y., Lopez, R., & Hunter, S. (2014). InterProScan 5: genome-scale protein function classification. Bioinformatics (Oxford, England), 30(9), 1236–1240.

Jumper, J., Evans, R., Pritzel, A., Green, T., Figurnov, M., Ronneberger, O., Tunyasuvunakool, K., Bates, R., Žídek, A., Potapenko, A., Bridgland, A., Meyer, C., Kohl, S. A. A., Ballard, A. J., Cowie, A., Romera-Paredes, B., Nikolov, S., Jain, R., Adler, J., … Hassabis, D. (2021). Highly accurate protein structure prediction with AlphaFold. Nature, 596(7873), 583–589.

Käll, L., Krogh, A., & Sonnhammer, E. L. L. (2004). A combined transmembrane topology and signal peptide prediction method. Journal of Molecular Biology, 338(5), 1027–1036.

Kanehisa, M., Furumichi, M., Sato, Y., Kawashima, M., & Ishiguro-Watanabe, M. (2023). KEGG for taxonomy-based analysis of pathways and genomes. Nucleic Acids Research, 51(D1), D587–D592.

Kanehisa, M., Sato, Y., & Morishima, K. (2016). BlastKOALA and GhostKOALA: KEGG tools for functional characterization of genome and metagenome sequences. Journal of Molecular Biology, 428(4), 726–731.

Kato, S., Ogasawara, A., Itoh, T., Sakai, H. D., Shimizu, M., Yuki, M., Kaneko, M., Takashina, T., & Ohkuma, M. (2022). Nanobdella aerobiophila gen. nov., sp. nov., a thermoacidophilic, obligate ectosymbiotic archaeon, and proposal of Nanobdellaceae fam. nov., Nanobdellales ord. nov. and Nanobdellia class. nov. International Journal of Systematic and Evolutionary Microbiology, 72(8). 10.1099/ijsem.0.005489

Katoh, K., & Standley, D. M. (2013). MAFFT multiple sequence alignment software version 7: improvements in performance and usability. Molecular Biology and Evolution, 30(4), 772–780.

Kellner, S., Spang, A., Offre, P., Szöllösi, G. J., Petitjean, C., & Williams, T. A. (2018). Genome size evolution in the Archaea. Emerging Topics in Life Sciences, 2(4), 595–605.

Kizina, J., Jordan, S. F. A., Martens, G. A., Lonsing, A., Probian, C., Kolovou, A., Santarella-Mellwig, R., Rhiel, E., Littmann, S., Markert, S., Stüber, K., Richter, M., Schweder, T., & Harder, J. (2022). Methanosaeta and “Candidatus Velamenicoccus archaeovorus.” Applied and Environmental Microbiology, 88(7), e0240721.

Kolmogorov, M., Yuan, J., Lin, Y., & Pevzner, P. A. (2019). Assembly of long, error-prone reads using repeat graphs. Nature Biotechnology, 37 (5), 540–546.

Krause, S., Gfrerer, S., von Kügelgen, A., Reuse, C., Dombrowski, N., Villanueva, L., Bunk, B., Spröer, C., Neu, T. R., Kuhlicke, U., Schmidt-Hohagen, K., Hiller, K., Bharat, T. A. M., Rachel, R., Spang, A., & Gescher, J. (2022). The importance of biofilm formation for cultivation of a Micrarchaeon and its interactions with its Thermoplasmatales host. Nature Communications, 13(1), 1735.

Krogh, A., Larsson, B., von Heijne, G., & Sonnhammer, E. L. (2001). Predicting transmembrane protein topology with a hidden Markov model: application to complete genomes. Journal of Molecular Biology, 305(3), 567–580.

Kuroda, K., Yamamoto, K., Nakai, R., Hirakata, Y., Kubota, K., Nobu, M. K., & Narihiro, T. (2022). Symbiosis between Candidatus Patescibacteria and Archaea discovered in wastewater-treating bioreactors. MBio, 13(5), e0171122.

La Cono, V., Messina, E., Rohde, M., Arcadi, E., Ciordia, S., Crisafi, F., Denaro, R., Ferrer, M., Giuliano, L., Golyshin, P. N., Golyshina, O. V., Hallsworth, J. E., La Spada, G., Mena, M. C., Merkel, A. Y., Shevchenko, M. A., Smedile, F., Sorokin, D. Y., Toshchakov, S. V., & Yakimov, M. M. (2020). Symbiosis between nanohaloarchaeon and haloarchaeon is based on utilization of different polysaccharides. Proceedings of the National Academy of Sciences of the United States of America, 117 (33), 20223–20234.

Lartillot, N., Rodrigue, N., Stubbs, D., & Richer, J. (2013). PhyloBayes MPI: phylogenetic reconstruction with infinite mixtures of profiles in a parallel environment. Systematic Biology, 62(4), 611–615.

Li, D., Liu, C.-M., Luo, R., Sadakane, K., & Lam, T.-W. (2015). MEGAHIT: an ultra-fast single-node solution for large and complex metagenomics assembly via succinct de Bruijn graph. Bioinformatics (Oxford, England), 31(10), 1674–1676.

McCutcheon, J. P., & Moran, N. A. (2011). Extreme genome reduction in symbiotic bacteria. Nature Reviews. Microbiology, 10(1), 13–26.

Mendler, K., Chen, H., Parks, D. H., Lobb, B., Hug, L. A., & Doxey, A. C. (2019). AnnoTree: visualization and exploration of a functionally annotated microbial tree of life. Nucleic Acids Research, 47 (9), 4442–4448.

Mikheenko, A., Saveliev, V., Hirsch, P., & Gurevich, A. (2023). WebQUAST: online evaluation of genome assemblies. Nucleic Acids Research, 51(W1), W601–W606.

Minh, B. Q., Schmidt, H. A., Chernomor, O., Schrempf, D., Woodhams, M. D., von Haeseler, A., & Lanfear, R. (2020). IQ-TREE 2: New models and efficient methods for phylogenetic inference in the genomic era. Molecular Biology and Evolution, 37 (5), 1530–1534.

Moissl-Eichinger, C., & Huber, H. (2011). Archaeal symbionts and parasites. Current Opinion in Microbiology, 14(3), 364–370.

Moreno, J., Nielsen, H., Winther, O., & Teufel, F. (2024). Predicting the subcellular location of prokaryotic proteins with DeepLocPro. Bioinformatics (Oxford, England), 40(12). 10.1093/bioinformatics/btae677

Nakayama, T., Harada, R., Yabuki, A., Nomura, M., Shiba, K., Inaba, K., & Inagaki, Y. (2025). Drastic genome reduction driven by parasitic lifestyle: Two complete genomes of endosymbiotic bacteria possibly hosted by a dinoflagellate. In bioRxiv. 10.1101/2025.01.03.631278

Nakayama, T., Nomura, M., Takano, Y., Tanifuji, G., Shiba, K., Inaba, K., Inagaki, Y., & Kawata, M. (2019). Single-cell genomics unveiled a cryptic cyanobacterial lineage with a worldwide distribution hidden by a dinoflagellate host. Proceedings of the National Academy of Sciences of the United States of America, 116(32), 15973–15978.

Nakayama, T., Nomura, M., Yabuki, A., Shiba, K., Inaba, K., & Inagaki, Y. (2024). Convergent reductive evolution of cyanobacteria in symbiosis with Dinophysiales dinoflagellates. Scientific Reports, 14(1), 12774.

Nishimura, Y., & Yoshizawa, S. (2022). The OceanDNA MAG catalog contains over 50,000 prokaryotic genomes originated from various marine environments. Scientific Data, 9(1), 305.

Nurk, S., Meleshko, D., Korobeynikov, A., & Pevzner, P. A. (2017). metaSPAdes: a new versatile metagenomic assembler. Genome Research, 27 (5), 824–834.

Parks, D. H., Chuvochina, M., Rinke, C., Mussig, A. J., Chaumeil, P.-A., & Hugenholtz, P. (2022). GTDB: an ongoing census of bacterial and archaeal diversity through a phylogenetically consistent, rank normalized and complete genome-based taxonomy. Nucleic Acids Research, 50(D1), D785–D794.

Pesant, S., Not, F., Picheral, M., Kandels-Lewis, S., Le Bescot, N., Gorsky, G., Iudicone, D., Karsenti, E., Speich, S., Troublé, R., Dimier, C., Searson, S., & Tara Oceans Consortium Coordinators. (2015). Open science resources for the discovery and analysis of Tara Oceans data. Scientific Data, 2(1), 150023.

Prjibelski, A., Antipov, D., Meleshko, D., Lapidus, A., & Korobeynikov, A. (2020). Using SPAdes de novo assembler. Current Protocols in Bioinformatics, 70(1), e102.

Quang, L. S., Gascuel, O., & Lartillot, N. (2008). Empirical profile mixture models for phylogenetic reconstruction. Bioinformatics (Oxford, England), 24(20), 2317–2323.

Read, T. D., Brunham, R. C., Shen, C., Gill, S. R., Heidelberg, J. F., White, O., Hickey, E. K., Peterson, J., Utterback, T., Berry, K., Bass, S., Linher, K., Weidman, J., Khouri, H., Craven, B., Bowman, C., Dodson, R., Gwinn, M., Nelson, W., … Fraser, C. M. (2000). Genome sequences of Chlamydia trachomatis MoPn and Chlamydia pneumoniae AR39. Nucleic Acids Research, 28(6), 1397–1406.

Reva, O., Messina, E., La Cono, V., Crisafi, F., Smedile, F., La Spada, G., Marturano, L., Selivanova, E. A., Rohde, M., Krupovic, M., & Yakimov, M. M. (2023). Functional diversity of nanohaloarchaea within xylan-degrading consortia. Frontiers in Microbiology, 14, 1182464.

Rodrigues-Oliveira, T., Wollweber, F., Ponce-Toledo, R. I., Xu, J., Rittmann, S. K.-M. R., Klingl, A., Pilhofer, M., & Schleper, C. (2023). Actin cytoskeleton and complex cell architecture in an Asgard archaeon. Nature, 613(7943), 332–339.

Sakai, H. D., Nur, N., Kato, S., Yuki, M., Shimizu, M., Itoh, T., Ohkuma, M., Suwanto, A., & Kurosawa, N. (2022). Insight into the symbiotic lifestyle of DPANN archaea revealed by cultivation and genome analyses. Proceedings of the National Academy of Sciences of the United States of America, 119(3), e2115449119.

Salazar, G., Paoli, L., Alberti, A., Huerta-Cepas, J., Ruscheweyh, H.-J., Cuenca, M., Field, C. M., Coelho, L. P., Cruaud, C., Engelen, S., Gregory, A. C., Labadie, K., Marec, C., Pelletier, E., Royo-Llonch, M., Roux, S., Sánchez, P., Uehara, H., Zayed, A. A., … Sunagawa, S. (2019). Gene expression changes and community turnover differentially shape the global ocean meta-transcriptome. Cell, 179(5), 1068-1083.e21.

Sayers, E. W., Beck, J., Bolton, E. E., Brister, J. R., Chan, J., Comeau, D. C., Connor, R., DiCuccio, M., Farrell, C. M., Feldgarden, M., Fine, A. M., Funk, K., Hatcher, E., Hoeppner, M., Kane, M., Kannan, S., Katz, K. S., Kelly, C., Klimke, W., … Sherry, S. T. (2024). Database resources of the National Center for Biotechnology Information. Nucleic Acids Research, 52(D1), D33–D43.

Seymour, C. O., Palmer, M., Becraft, E. D., Stepanauskas, R., Friel, A. D., Schulz, F., Woyke, T., Eloe-Fadrosh, E., Lai, D., Jiao, J.-Y., Hua, Z.-S., Liu, L., Lian, Z.-H., Li, W.-J., Chuvochina, M., Finley, B. K., Koch, B. J., Schwartz, E., Dijkstra, P., … Hedlund, B. P. (2023). Hyperactive nanobacteria with host-dependent traits pervade Omnitrophota. Nature Microbiology, 8(4), 727–744.

Shen, W., Sipos, B., & Zhao, L. (2024). SeqKit2: A Swiss army knife for sequence and alignment processing. IMeta, 3(3), e191.

Shigenobu, S., Watanabe, H., Hattori, M., Sakaki, Y., & Ishikawa, H. (2000). Genome sequence of the endocellular bacterial symbiont of aphids Buchnera sp. APS. Nature, 407 (6800), 81–86.

Shimodaira, H. (2002). An approximately unbiased test of phylogenetic tree selection. Systematic Biology, 51(3), 492–508.

Sørensen, M. E. S., Stiller, M. L., Kröninger, L., & Nowack, E. C. M. (2024). Protein import into bacterial endosymbionts and evolving organelles. The FEBS Journal. 10.1111/febs.17356

Spang, A., Saw, J. H., Jørgensen, S. L., Zaremba-Niedzwiedzka, K., Martijn, J., Lind, A. E., van Eijk, R., Schleper, C., Guy, L., & Ettema, T. J. G. (2015). Complex archaea that bridge the gap between prokaryotes and eukaryotes. Nature, 521(7551), 173–179.

St John, E., Liu, Y., Podar, M., Stott, M. B., Meneghin, J., Chen, Z., Lagutin, K., Mitchell, K., & Reysenbach, A.-L. (2019). A new symbiotic nanoarchaeote (Candidatus Nanoclepta minutus) and its host (Zestosphaera tikiterensis gen. nov., sp. nov.) from a New Zealand hot spring. Systematic and Applied Microbiology, 42(1), 94–106.

Sunagawa, S., Coelho, L. P., Chaffron, S., Kultima, J. R., Labadie, K., Salazar, G., Djahanschiri, B., Zeller, G., Mende, D. R., Alberti, A., Cornejo-Castillo, F. M., Costea, P. I., Cruaud, C., d’Ovidio, F., Engelen, S., Ferrera, I., Gasol, J. M., Guidi, L., Hildebrand, F., … Bork, P. (2015). Structure and function of the global ocean microbiome. Science (New York, N.Y.), 348(6237), 1261359.

Susko, E., & Roger, A. J. (2007). On reduced amino acid alphabets for phylogenetic inference. Molecular Biology and Evolution, 24(9), 2139–2150.

Tahon, G., Geesink, P., & Ettema, T. J. G. (2021). Expanding Archaeal diversity and phylogeny: Past, present, and future. Annual Review of Microbiology, 75(1), 359–381.

Tanizawa, Y., Fujisawa, T., & Nakamura, Y. (2018). DFAST: a flexible prokaryotic genome annotation pipeline for faster genome publication. Bioinformatics (Oxford, England), 34(6), 1037–1039.

Tully, B. J., Graham, E. D., & Heidelberg, J. F. (2018). The reconstruction of 2,631 draft metagenome-assembled genomes from the global oceans. Scientific Data, 5(1), 170203.

van Kempen, M., Kim, S. S., Tumescheit, C., Mirdita, M., Lee, J., Gilchrist, C. L. M., Söding, J., & Steinegger, M. (2024). Fast and accurate protein structure search with Foldseek. Nature Biotechnology, 42(2), 243–246.

van Wolferen, M., Pulschen, A. A., Baum, B., Gribaldo, S., & Albers, S.-V. (2022). The cell biology of archaea. Nature Microbiology, 7 (11), 1744–1755.

Vaser, R., & Šikić, M. (2021). Time- and memory-efficient genome assembly with Raven. Nature Computational Science, 1(5), 332–336.

Wang, H.-C., Li, K., Susko, E., & Roger, A. J. (2008). A class frequency mixture model that adjusts for site-specific amino acid frequencies and improves inference of protein phylogeny. BMC Evolutionary Biology, 8(1), 331.

Wang, H.-C., Minh, B. Q., Susko, E., & Roger, A. J. (2018). Modeling site heterogeneity with posterior mean site frequency profiles accelerates accurate phylogenomic estimation. Systematic Biology, 67 (2), 216–235.

Waters, E., Hohn, M. J., Ahel, I., Graham, D. E., Adams, M. D., Barnstead, M., Beeson, K. Y., Bibbs, L., Bolanos, R., Keller, M., Kretz, K., Lin, X., Mathur, E., Ni, J., Podar, M., Richardson, T., Sutton, G. G., Simon, M., Soll, D., … Noordewier, M. (2003). The genome of Nanoarchaeum equitans: insights into early archaeal evolution and derived parasitism. Proceedings of the National Academy of Sciences of the United States of America, 100(22), 12984–12988.

Wick, R. R., Judd, L. M., Gorrie, C. L., & Holt, K. E. (2017). Unicycler: Resolving bacterial genome assemblies from short and long sequencing reads. PLoS Computational Biology, 13(6), e1005595.

Wrede, C., Dreier, A., Kokoschka, S., & Hoppert, M. (2012). Archaea in symbioses. Archaea (Vancouver, B.C.), 2012, 596846.

Wurch, L., Giannone, R. J., Belisle, B. S., Swift, C., Utturkar, S., Hettich, R. L., Reysenbach, A.-L., & Podar, M. (2016). Genomics-informed isolation and characterization of a symbiotic Nanoarchaeota system from a terrestrial geothermal environment. Nature Communications, 7, 12115.

Zaremba-Niedzwiedzka, K., Caceres, E. F., Saw, J. H., Bäckström, D., Juzokaite, L., Vancaester, E., Seitz, K. W., Anantharaman, K., Starnawski, P., Kjeldsen, K. U., Stott, M. B., Nunoura, T., Banfield, J. F., Schramm, A., Baker, B. J., Spang, A., & Ettema, T. J. G. (2017). Asgard archaea illuminate the origin of eukaryotic cellular complexity. Nature, 541(7637), 353–358.

Zhu, Q., Mai, U., Pfeiffer, W., Janssen, S., Asnicar, F., Sanders, J. G., Belda-Ferre, P., Al-Ghalith, G. A., Kopylova, E., McDonald, D., Kosciolek, T., Yin, J. B., Huang, S., Salam, N., Jiao, J.-Y., Wu, Z., Xu, Z. Z., Cantrell, K., Yang, Y., … Knight, R. (2019). Phylogenomics of 10,575 genomes reveals evolutionary proximity between domains Bacteria and Archaea. Nature Communications, 10(1), 5477.

