## Supplementary Figure S1 for "A cellular entity retaining only its replicative core: Hidden archaeal lineage with an ultra-reduced genome"

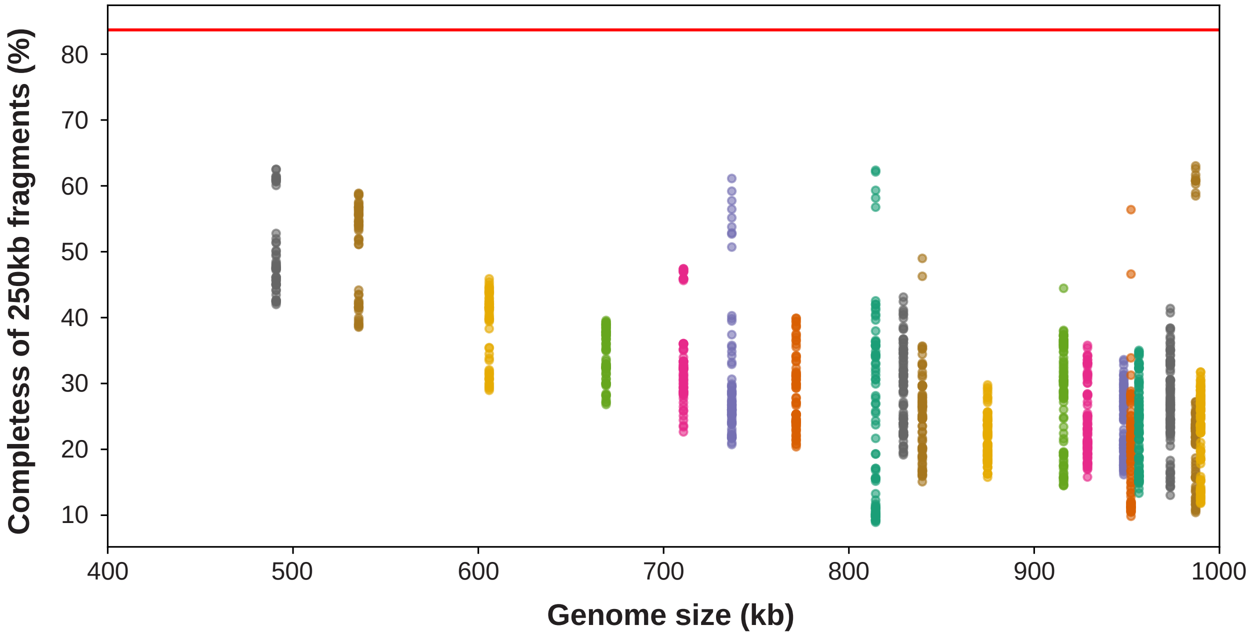


**Figure S1.** CheckM2 completeness comparison of 250 kb archaeal genome fragments. Scatter plots indicate the completeness of 250 kb genome fragments calculated by CheckM2 (y-axis) and the genome sizes of original genomes (x-axis). The dozens of the fragments were generated from the same original genome by the 10 kb-sliding windows as dots of the same color apppeares in a vertical line. The archeal genomes comprising one contig, smaller than 1 Mbp, and selected as 'representative' in GTDB are shown. A red horizontal line represents the completeness of the Sukunaarchaeum genome (83.68%).
