## Supplementary Figure S2 for "A cellular entity retaining only its replicative core: Hidden archaeal lineage with an ultra-reduced genome"

### A. Ribosomal protein L23 (SKNCREG01_0030)

```
Sukunaarchaeum     Sukunaarchaeum     SKNSIEKISLSVKSKALLDHCKTITFYFKNNLSLTKNQIKNQILALYQLRILK
Methanobacteriati  GCA_000091665.1    AFDVIKAPVVTEKTVRMIEEENKLVFYVDRR--ATKQDIKRAMKELFDVEVEK
Methanobacteriati  GCA_023261745.1    PFEIIIKPYVTEKTMFMIEKENKLAFIVKRD--ATKMDIKWAVERAFGVKVES
Methanobacteriati  GCA_018609935.1    --MVIKKPHITEKAMDLMDFENKLQFIVEDK--ATKKEIKQEVEDQWGEEVEL
Thermoproteati     GCA_022839705.1    PSDIIIRPVSSESALEKVEKENKILFIVDPK--SNKIQIKQAVEKLYGVKVKS
Promethearchaeati  GCA_016294985.1    YYQVIHNVVISEKLFDMIENENKLAFIVERR--ANKREIKKAVELLYEVKVDR
Nanobdellati       GCA_902384795.1    -MALIKYPITTEKAIGMIELQNKIVFVVDKS--ATKPGVKEEVEKTFGVKVAQ
Nanobdellati       GCA_018304445.1    NFEIIKFALISEKAVNMIERENKLTFIVNQN--ADKPSIKKAIEEMHKVKVDN

                                      ISKIYDFKRRKLKALIRYADESLPTELQWVKE-H
                                      VNTLITPK-GEKKAYVKLKEGYDASKIAASLGIY
                                      VNVLYSRK-G-KKAFVKLRPEFNADEIAVRIGIF
                                      VNTMITPS-GEKKATVKFKEEGISDDIASRMGMF
                                      VNTMITPS-AEKKAAVVLEKEFSATELATKIGLL
                                      VNTLILCD-GRKKAYVRLSDKDDASDLAAKLGMF
                                      VNILITPK-GQKRAYVKLAKGFSADEVAAKMGVA
                                      VKIIRDRK-GRKKAIVKIGKGFKASDVATRLGVI
```

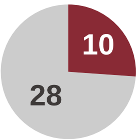


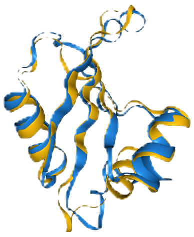

pident: 16.0%
evalue: 6.08e-4
RMSD: 2.71

### B. Ribosomal protein S10 (SKNCREG01_0740)

```
Sukunaarchaeum     Sukunaarchaeum     -MQ-LKFLVRSNSFKDLKRLLEELK--------DVT-IVHFPRKDIKFVLPKN
Methanobacteriati  GCA_000091665.1    -MQRARIKLSSTDHKVLDEICRQIKEIAEKTGVDISGPIPLPTKVLRVVTRK-
Methanobacteriati  GCA_023261745.1    MVYKARIRLSGNDYRKVEEVCDEIKNIAERTGVEIHGPIPLPTKRLIVPVRK-
Methanobacteriati  GCA_018609935.1    -MQKARVRLSGTDHEELDEVADEITEIAERTGVEFSGPVPLPTKDLEVPIRK-
Thermoproteati     GCA_022839705.1    MPQKARIMLSSTDFQKLEDVCTQIRKIAEKTGVKIAGPVPLPTKKLLVPTLK-
Promethearchaeati  GCA_016294985.1    MVQRARIRLLGKDTTELQNVCNQLKEIAQKTGARFNGPIPLPTKKIKIPVRK-
Nanobdellati       GCA_902384795.1    -MGKARIKLVGLNPKELEGVCQQIIEIVNTTGVEKRGPIPLPTRKLRVPTRR-
Nanobdellati       GCA_018304445.1    -MQKARIKLSSPDYDTLNGICTQIIDVAERTGVKHSGSIPLPTKKMVIPTRK-

                                      YGIKGCRGIGTYDRIEYNFWKKIITVDISKIGIIGNF---IAEGLDIDLIED
                                      --SPDGEGSSTFDRWTMKIHKRLIDIDADERALRHIMKIRIPDNVQIEIQFK
                                      --TPCGDGSATWDHWEMRIHKRLIDIEADERALRQLMRIQIPDGIHIEIELK
                                      --SPDGEGNATWDHWEMRVHKRLIDLEANERALRQLMRVQVPQEVSIEIELE
                                      --SPAGEGTKTWDKWEMRIHKRLIDVDADDRTMRQMMRIQVPDDVFIEIELV
                                      --TPCGEGTPTWEQWQMRVHKRIVILDANERTMSQIMRIRVPDSIYIEIELV
                                      --TPCGDGSDTYEAWEMRVHKRVIEVAGDERTLRQIMRVKVPDTVHIEISLH
                                      --SPCGGGTHTFDHWEMRVHKRLIDIEADERTLRRIMRVEIPNDEGVLVELP
```

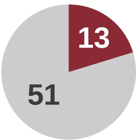


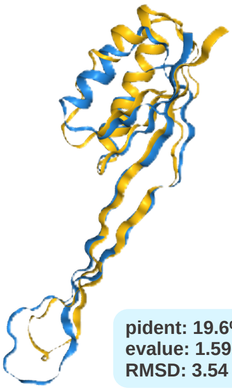

pident: 19.6%
evalue: 1.59e-5
RMSD: 3.54

### C. Translation initiation factor 2 beta (SKNCREG01_0720)

```
Sukunaarchaeum     Sukunaarchaeum     MT------------IFEDLLNSIGKDH--QEQSGL-KLELTVEKQGLIKIKNL
Methanobacteriati  GCA_000091665.1    MSDLENIDYYDYKALLKRARSQIPDYVFQKDRFELPEIEILIEG-NRTIIRNF
Methanobacteriati  GCA_023261745.1    MT-----DIFDYDSLLKRAMALKPKEVDSGERFQIPDAEIIIEG-KNTILKNF
Methanobacteriati  GCA_018609935.1    M---------EYDEALDKAYENITEVTKHGERFEPPEFNIRVEG-NSTIITNF
Thermoproteati     GCA_022839705.1    MQ-------MDYESLLKRGMSRIPEKAIKSSKYETPQTESNVIG-AKTVVYNF
Promethearchaeati  GCA_016294985.1    MD----VNFFNYDEMLKKAKEQVPPDIYHHKRFEIPVVDLFIEG-NRTVVQNW
Nanobdellati       GCA_902384795.1    MD--------DYEKLLDKAYANLPKKVLTQERFEMPAVDSFIQG-TKTIVKNF
Nanobdellati       GCA_018304445.1    ME------QAEYEKLLDRAYAALPKEALTKERFEMPVADSFIQG-QKTNVKNF

                                      KEILK-FFEDKTSEWFFYQFSRKCG--ITFDKNYR-FKTIIDQKNLEKKLIEL
                                      RELAKAVNRD--EEFFAKYLLKETGSAGNLEGGRLILQRRISPELLKSRINDF
                                      KEFCDRLDRP--PEHFSKFLFRELGTAGQLQGERLIFKGKVSSLDIKNRINDY
                                      RDVAEKLDRD--ENHLMKHVLGEIGTAGHLENRRARLKGEFSEENLQEALKTY
                                      KDVAAKLNRD--PNHLLKFMVRELATSGVLEDARASLHGRFMKESLDELVVRY
                                      KDITTRLNRD--PAHLLKFLTRELATAGTMDGPRAIFQGKFSKNFLNDLIERY
                                      DFVCQKLRRD--PNQVAKYLYKELAAPGSVQGQRLMLQGKFGDKMLNEKLENY
                                      GSILKTIRRE--PEHMLKYITKEIGTQATMQEDRLVLNGKFTSKQVNDIFTNY

                                      YYKYQVCSSCKKNLV---KTT-DM-QKICLSCGFIEKILQDS
                                      LREYVICRECGKPDTKIIKEG-RVHLLKCMACGAIRPIRMI-
                                      IKTYVLCYECGSPDTILKKEG-RVEILVCKACGAIRPVNAKI
                                      IEKYVLCSECNRPDTRLEKQK-GTLTLKCEACGAFSPVK---
                                      IKAYVLCTSCNKPDTKLLRED-RMTFIVCEVCGAKNPAKTLV
                                      AKRYLICPTCTSPDTILLKEG-PYHFIKCEACGAKEAVPKLK
                                      AKTYVLCKECGKPDTNLIEEGNRIHTLVCEACGARAPARGV-
                                      IKMYVLCHECLRPDTKIADEH-GVKLLKCEACGASSPVKRL-
```

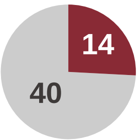


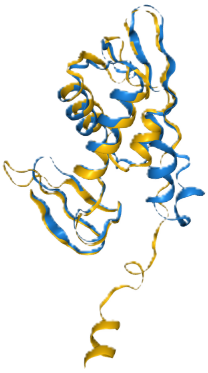

pident: 17.9%
evalue: 4.09e-5
RMSD: 8.95

**Figure S2.** Alignments of primary sequences and tertiary structures of genes annotated by Foldseek. Three examples of Sukunaarchaeum genes that were not annotated by the homology search based on amino acid sequences but were annotated in the AlphaFold/Foldseek analysis are shown. (A) Ribosomal protein L23. (B) Ribosomal protein S10. (C) Translation initiation factor 2 beta. The amino acid sequence alignments show homologs of Sukunaarchaeum and seven diverse archaea species, and highly conserved sites with five or more species having the same amino acids are shown in red. The pie charts show the proportion of highly conserved sites in which Sukunaarchaeum has a conserved amino acid. The right panels show the tertiary structural aignments of Sukunaarchaeum in blue and the best hit homolog in yellow. The identity, E-value, and RMSD of the Foldseek hits are shown at the bottom right.
