## Supplementary Figure S3 for "A cellular entity retaining only its replicative core: Hidden archaeal lineage with an ultra-reduced genome"

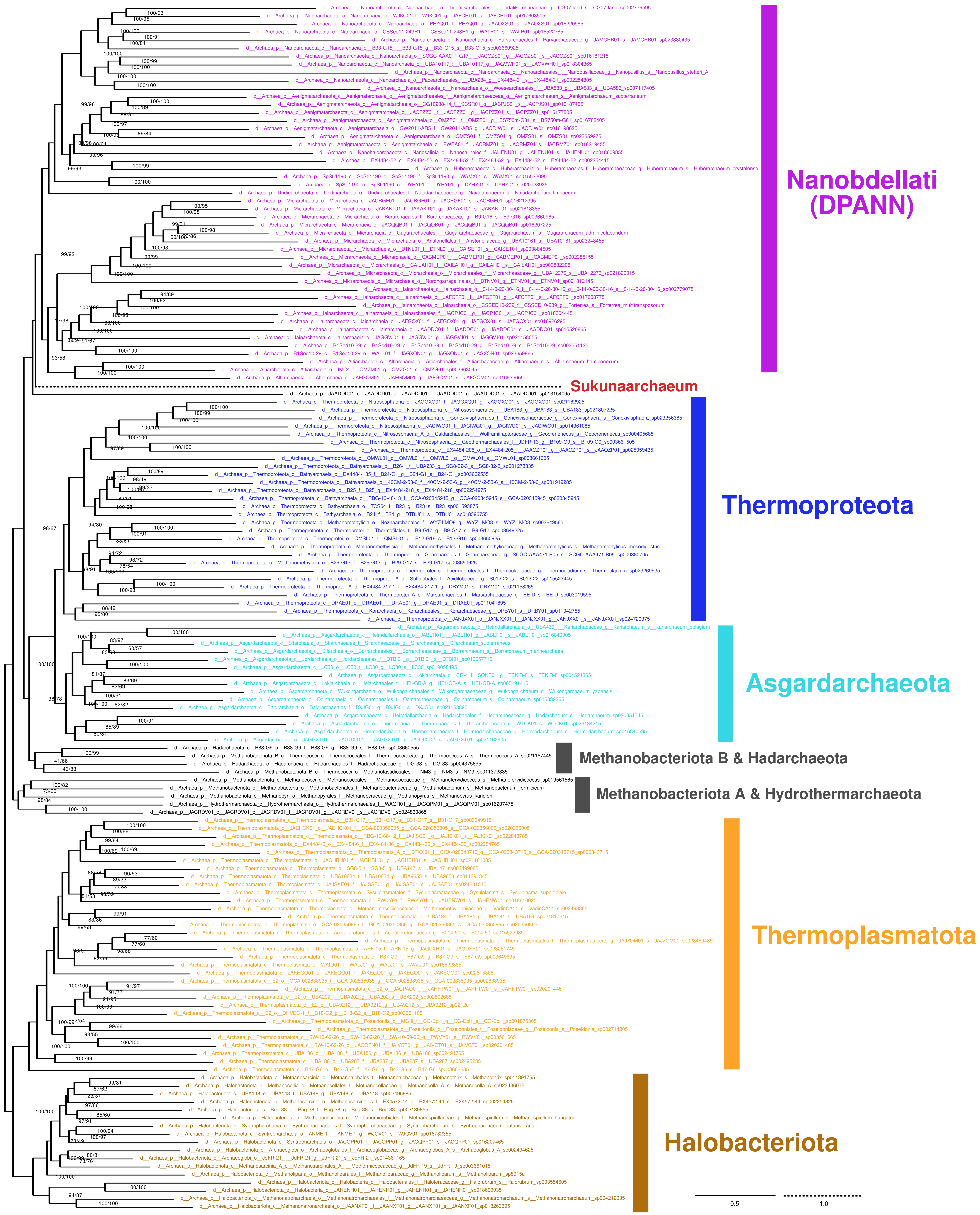


**Figure S3.** Maximum likelihood (ML) phylogenetic tree based on 70 gene alignment. The tree was inferred using IQ-Tree under the LG+C60+F+I+R10 model. The statistical support for each bipartition in the ML tree was calculated by 1000-replicate ultrafast bootstrap approximation (left) and 100-replicate nonparametric bootstrap analysis (right) with the posterior mean site frequency (PMSF) modeling. The labeling of leaves follows the taxonomy of GTDB r214.
