## Supplementary Figure S4 for "A cellular entity retaining only its replicative core: Hidden archaeal lineage with an ultra-reduced genome"

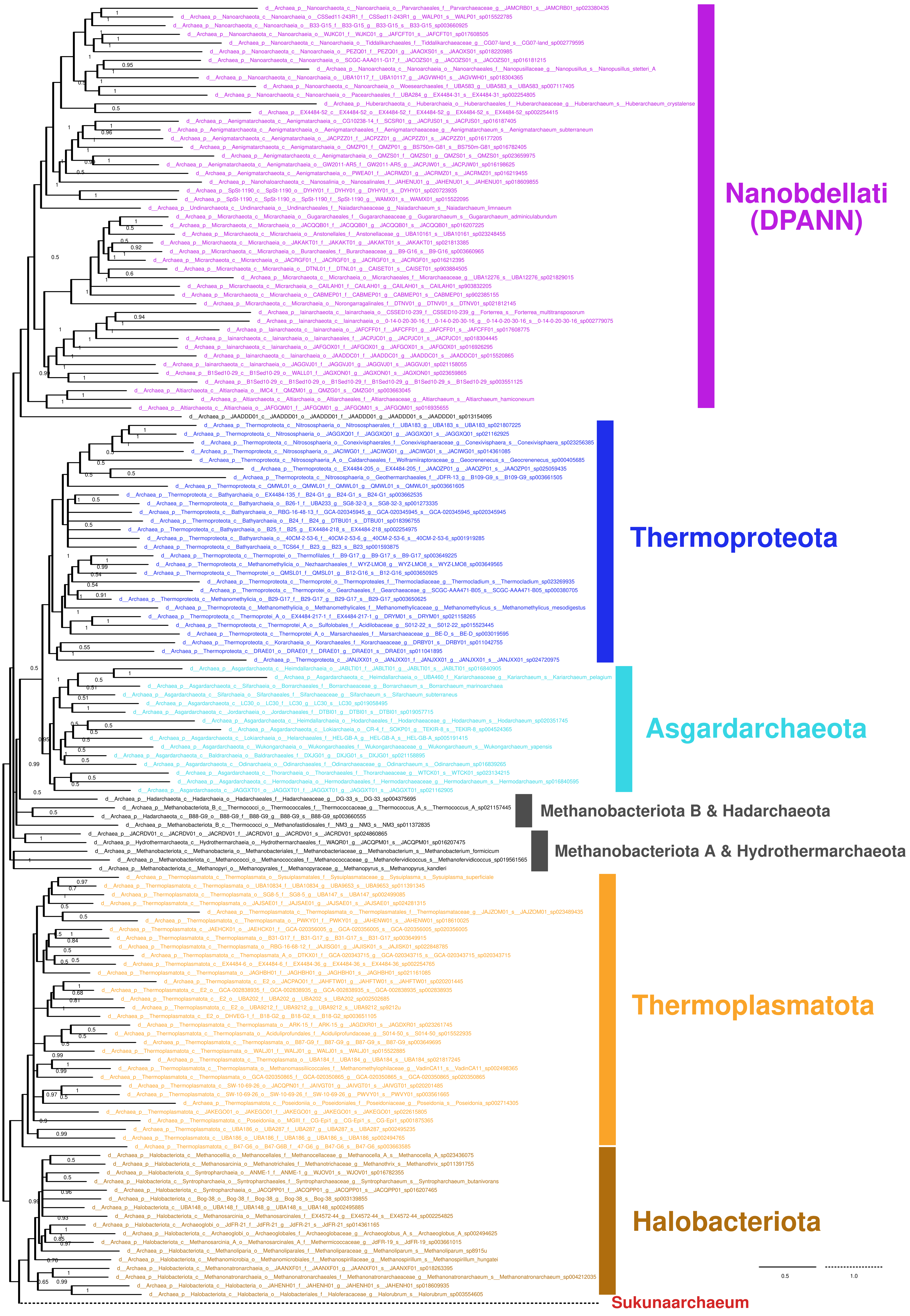


**Figure S4.** Bayesian phylogenetic tree based on 70 gene alignment.
The tree was inferred using PhyloBayes under the GTR+CAT+Γ4 model. The statistical support for each bipartition in the consensus tree was Bayesian posterior probability. The labeling of leaves follows the taxonomy of GTDB r214.
