## Supplementary Figure S5 for "A cellular entity retaining only its replicative core: Hidden archaeal lineage with an ultra-reduced genome"

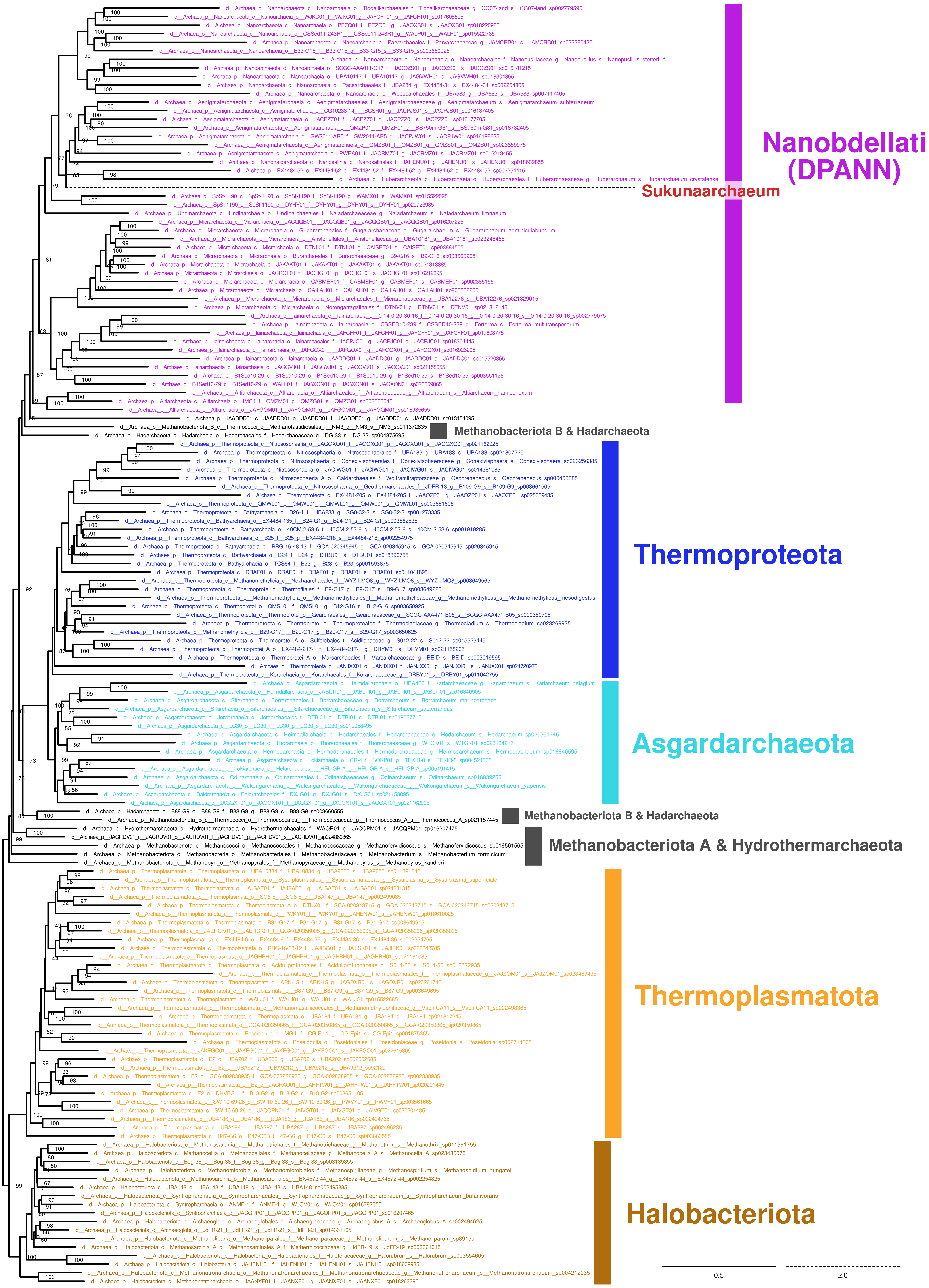


**Figure S5.** Maximum likelihood (ML) phylogenetic tree based on SR4 recoded 70 gene alignment. The tree was inferred using IQ-Tree under the GTR+SR4C60+F+I+R8 model. The statistical support for each bipartition in the ML tree was calculated by 1000-replicate ultrafast bootstrap approximation. The labeling of leaves follows the taxonomy of GTDB r214.
