## Supplementary Figure S7 for "A cellular entity retaining only its replicative core: Hidden archaeal lineage with an ultra-reduced genome"

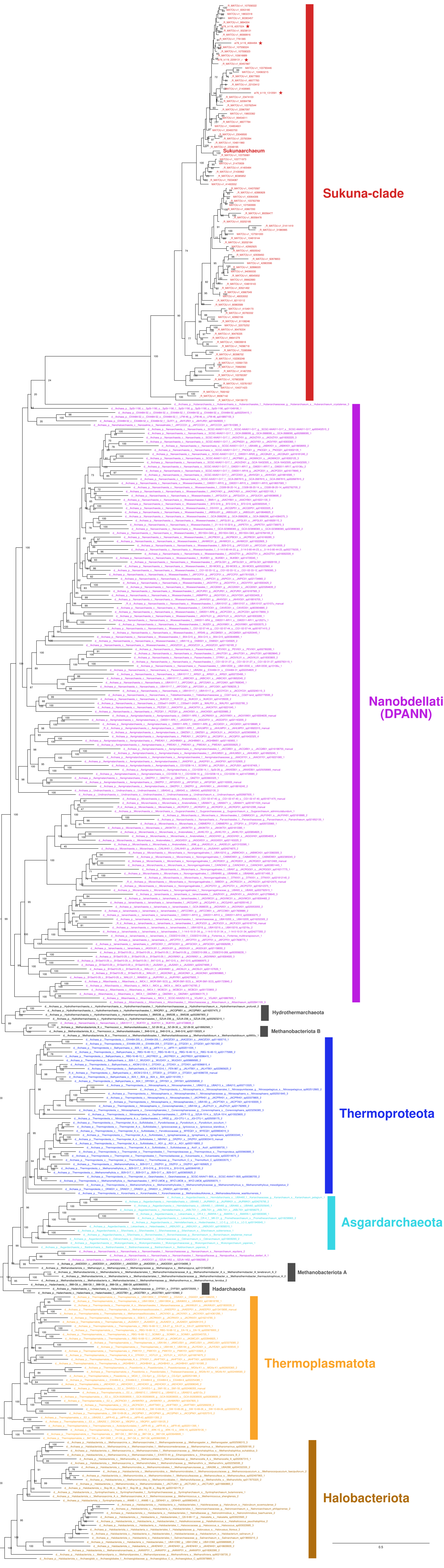


**Figure S7.** Maximum likelihood (ML) phylogenetic tree based on 23S rRNA gene alignment. The tree was inferred using IQ-Tree under the GTR+F+I+G4 model. The statistical support for each bipartition in the ML tree was calculated by 1000-replicate ultrafast bootstrap approximation. The labeling of leaves follows the taxonomy of GTDB r214. The stars represent the sequences detected from metagenomic assembly of *Tara* Oceans sampling station 76.
