## Supplementary Figure S8 for "A cellular entity retaining only its replicative core: Hidden archaeal lineage with an ultra-reduced genome"

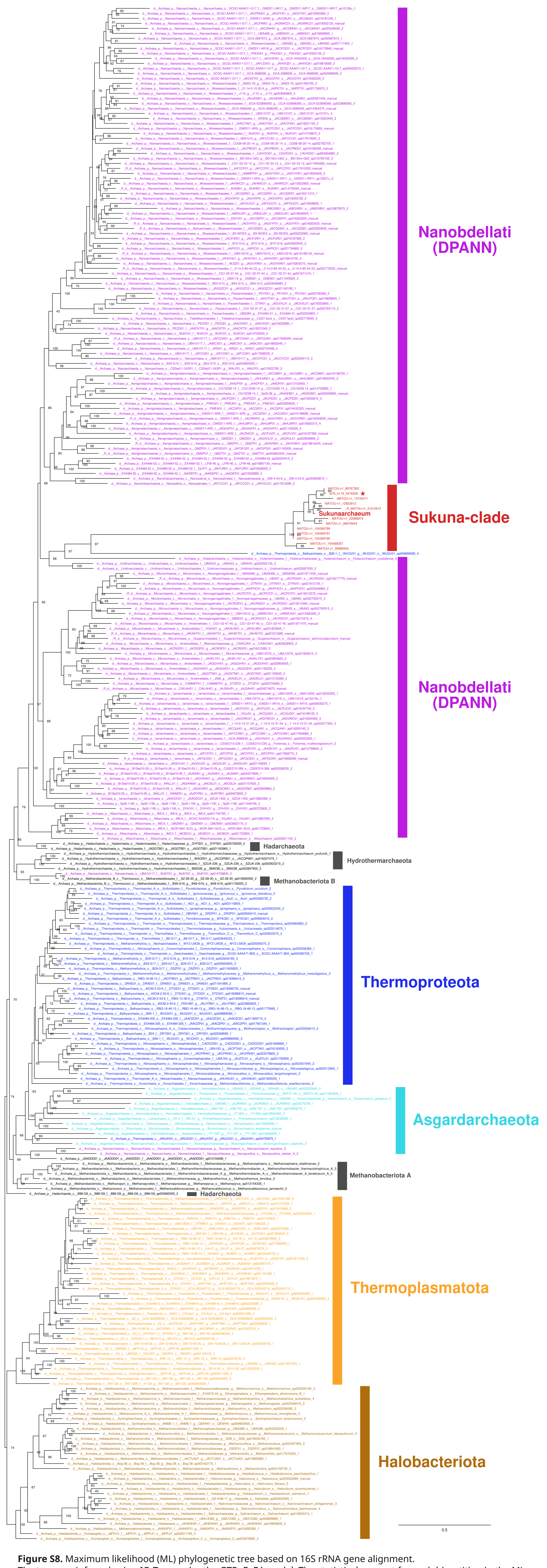


**Figure S8.** Maximum likelihood (ML) phylogenetic tree based on 16S rRNA gene alignment. The tree was inferred using IQ-Tree under the GTR+F+Γ4 model. The statistical support for each bipartition in the ML tree was calculated by 1000-replicate ultrafast bootstrap approximation. The labeling of leaves follows the taxonomy of GTDB r214. The star represents the sequence detected from metagenomic assembly of *Tara* Oceans sampling station 76.
