## Supplementary Figure S9 for "A cellular entity retaining only its replicative core: Hidden archaeal lineage with an ultra-reduced genome"

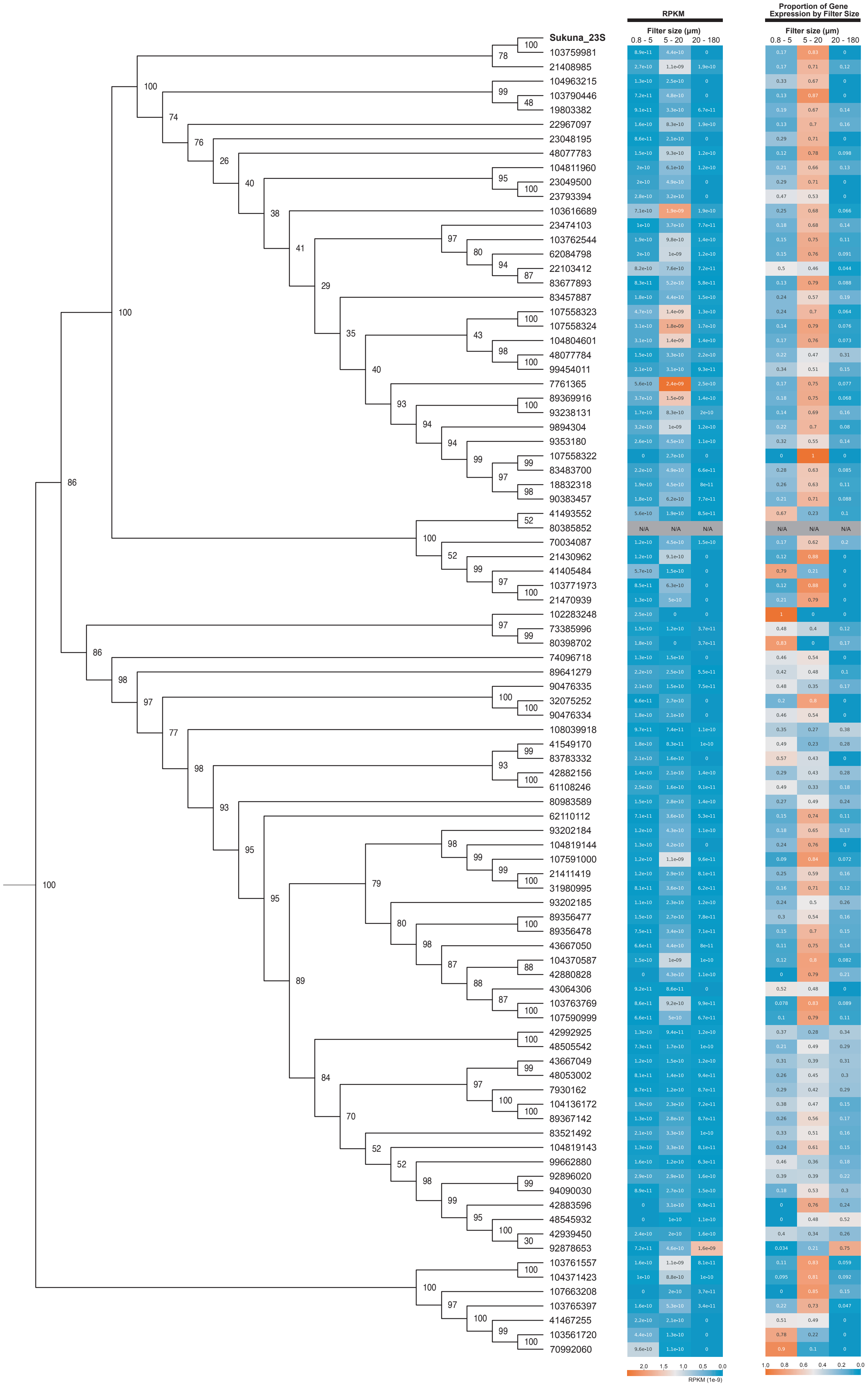


**Figure S9**. Relative abundance patterns across size fractions for Sukuna-clade 23S rRNA gene sequences derived from *Tara* Oceans metatranscriptome data. The cladogram on the left depicts the phylogenetic relationships among the 23S rRNA gene sequence from *Candidatus* Sukunaarchaeum mirabile (labelled "Sukuna_23S") and related sequences identified from the Marine Atlas of *Tara* Oceans Unigenes (MATOU) database. The central heatmap displays the average abundance (e.g., RPKM values) of each sequence within three distinct filter size fractions (0.8–5 μm, 5–20 μm, and 20–180 μm) from *Tara* Oceans samples. The right heatmap shows the relative abundance of each sequence normalized across these three size fractions (sum of relative abundances equals 1 for each sequence/row), illustrating the size fraction in which each sequence is proportionally most abundant.
