## Supplementary Figure S10 for "A cellular entity retaining only its replicative core: Hidden archaeal lineage with an ultra-reduced genome"

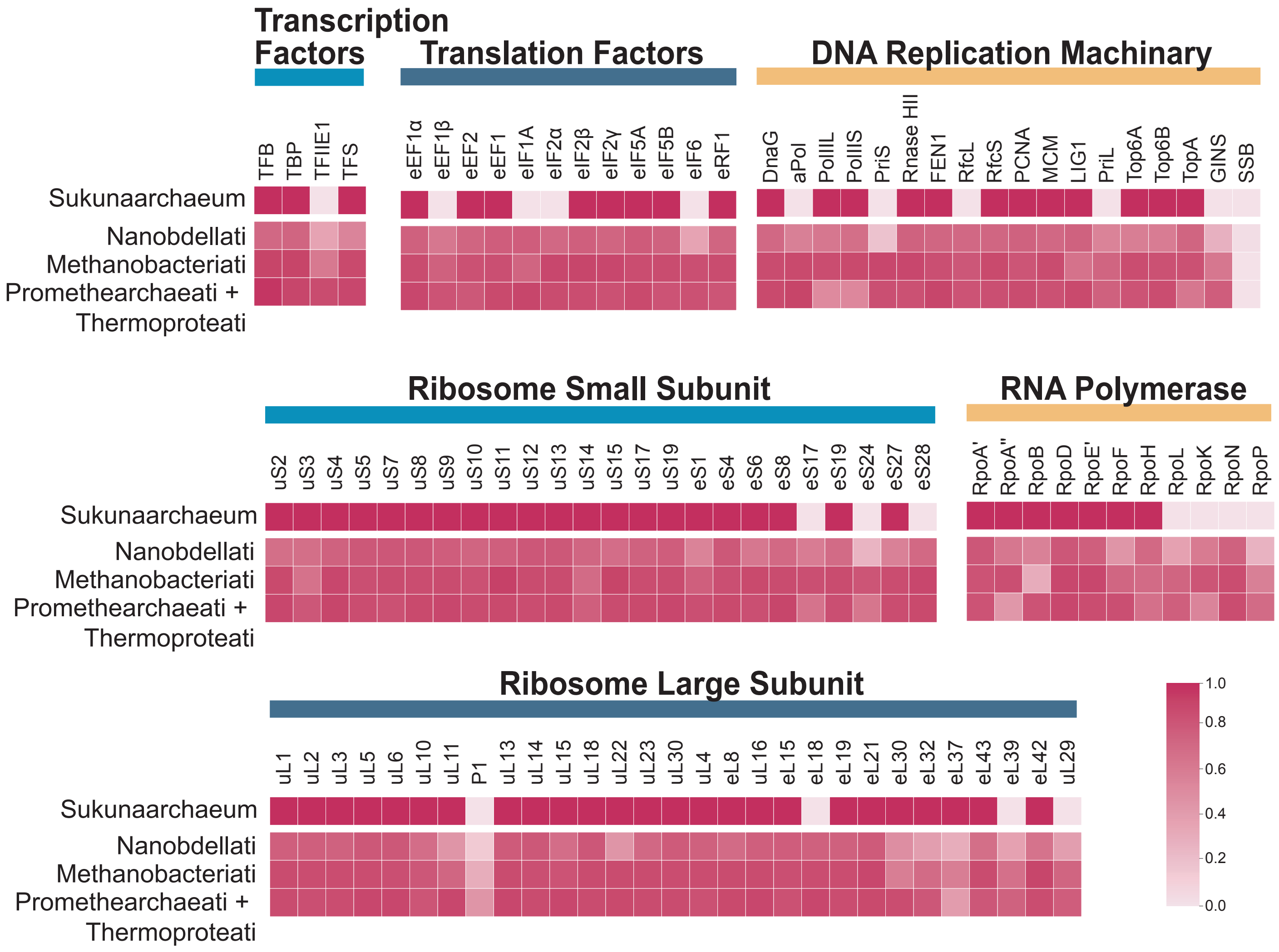


**Figure S10**. Conservation profile of core genetic information processing machinery genes across major archaeal lineages. Heatmap showing the proportion of genomes encoding homologs for core genes involved in transcription, translation, and DNA replication within *Ca.* Sukunaarchaeum mirabile and representative major archaeal groups (Nanobdellati, Methanobacteriati, and Promethearchaeati + Thermoproteati). Color intensity corresponds to the fraction of analyzed genomes within each group that possess a recognizable homolog of the respective gene.
