## Supplementary Figure S11 for "A cellular entity retaining only its replicative core: Hidden archaeal lineage with an ultra-reduced genome"

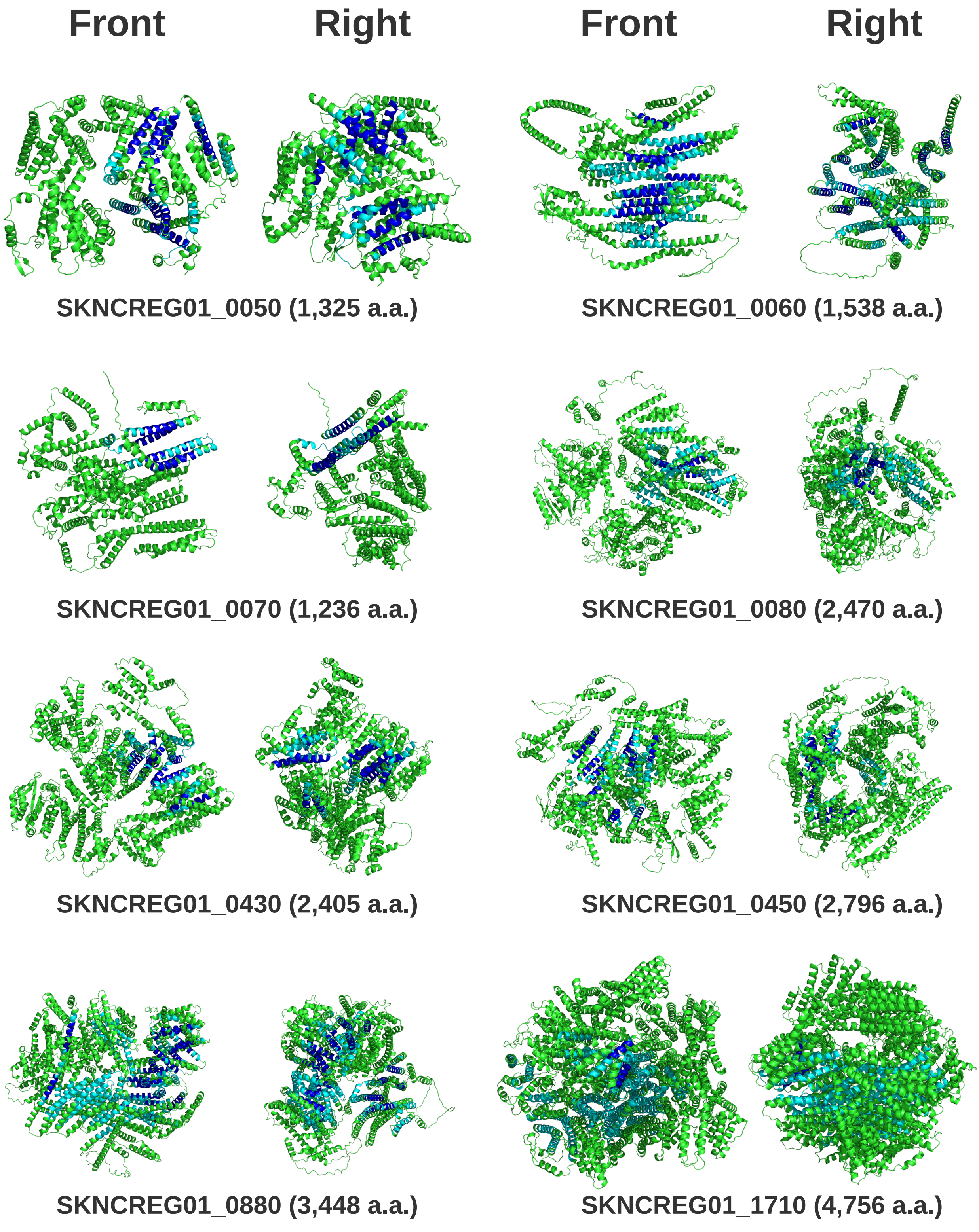


**Figure S11.** Predicted tertiary structures of eight hypothetical proteins of Sukunaarchaeum. Each structure is shown from the front and right views. Transmembrane (TM) domains were predicted by three *in silico* tools. TM regions predicted by all three tools are shown in blue, and TM regions predicted by at least one tool are shown in cyan.
