## Supplementary Figure S12 for "A cellular entity retaining only its replicative core: Hidden archaeal lineage with an ultra-reduced genome"

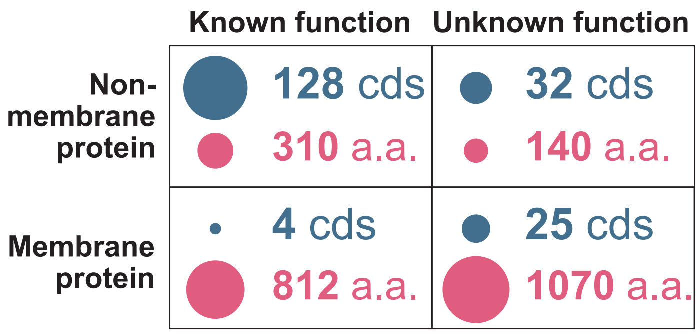


**Figure S12**. Numbers and average lengths of predicted membrane and non-membrane proteins in Sukunaarchaeum, categorized by known or unknown function.
